# Distinct CD8^+^ T-cell types Associated with COVID-19 Severity in Unvaccinated HLA-A2^+^ Patients

**DOI:** 10.1101/2025.01.12.632164

**Authors:** Kazuya Masuda, Sho Iketani, Lihong Liu, Jing Huang, Yujie Qiao, Jayesh Shah, Meredith L. McNairy, Christine Groso, Christopher Ricupero, Lucas F. Loffredo, Qian Wang, Lawrence Purpura, Jordana Grazziela Alves Coelho-dos-Reis, Zizhang Sheng, Michael T Yin, Moriya Tsuji

## Abstract

Although emerging data have revealed the critical role of memory CD8^+^ T cells in preventing and controlling SARS-CoV-2 infection, virus-specific CD8^+^ T-cell responses against SARS-CoV-2 and its memory and innate-like subsets in unvaccinated COVID-19 patients with various disease manifestations in an HLA-restricted fashion remain to be understood. Here, we show the strong association of protective cellular immunity with mild COVID-19 and unique cell types against SARS-CoV-2 virus in an HLA-A2 restricted manner. ELISpot assays reveal that SARS-CoV-2-specific CD8^+^ T-cell responses in mild COVID-19 patients are significantly higher than in severe patients, whereas neutralizing antibody responses against SARS-CoV-2 virus significantly correlate with disease severity. Single-cell analyses of HLA-A2-restricted CD8^+^ T cells, which recognize highly conserved immunodominant SARS-CoV-2-specific epitopes, demonstrate divergent profiles in unvaccinated patients with mild versus severe disease. CD8^+^ T-cell types including cytotoxic *KLRB1*^+^ CD8αα cells with innate-like T-cell signatures, *IFNG*^hi^*ID3*^hi^ memory cells and *IL7R*^+^ proliferative stem cell-like memory cells are preferentially observed in mild COVID-19, whereas distinct terminally-differentiated T-cell subsets are predominantly detected in severe COVID-19: highly activated *FASL*^hi^ T-cell subsets and early-terminated or dysfunctional *IL4R*^+^ *GATA3*^+^ stem cell-like memory T-cell subset. In conclusion, our findings suggest that unique and contrasting SARS-CoV-2-specific CD8^+^ T-cell profiles may dictate COVID-19 severity.

Graphical abstract.
SARS-CoV-2 epitope-specific CD8^+^ T-cell subtypes associated with mild or severe COVID-19 patients.
Potent memory CD8^+^ T-cell subtypes with gene signature in mild COVID-19 patients (upper) and dysfunctional CD8^+^ T-cell subtypes with gene signature in severe COVID-19 patients (lower).

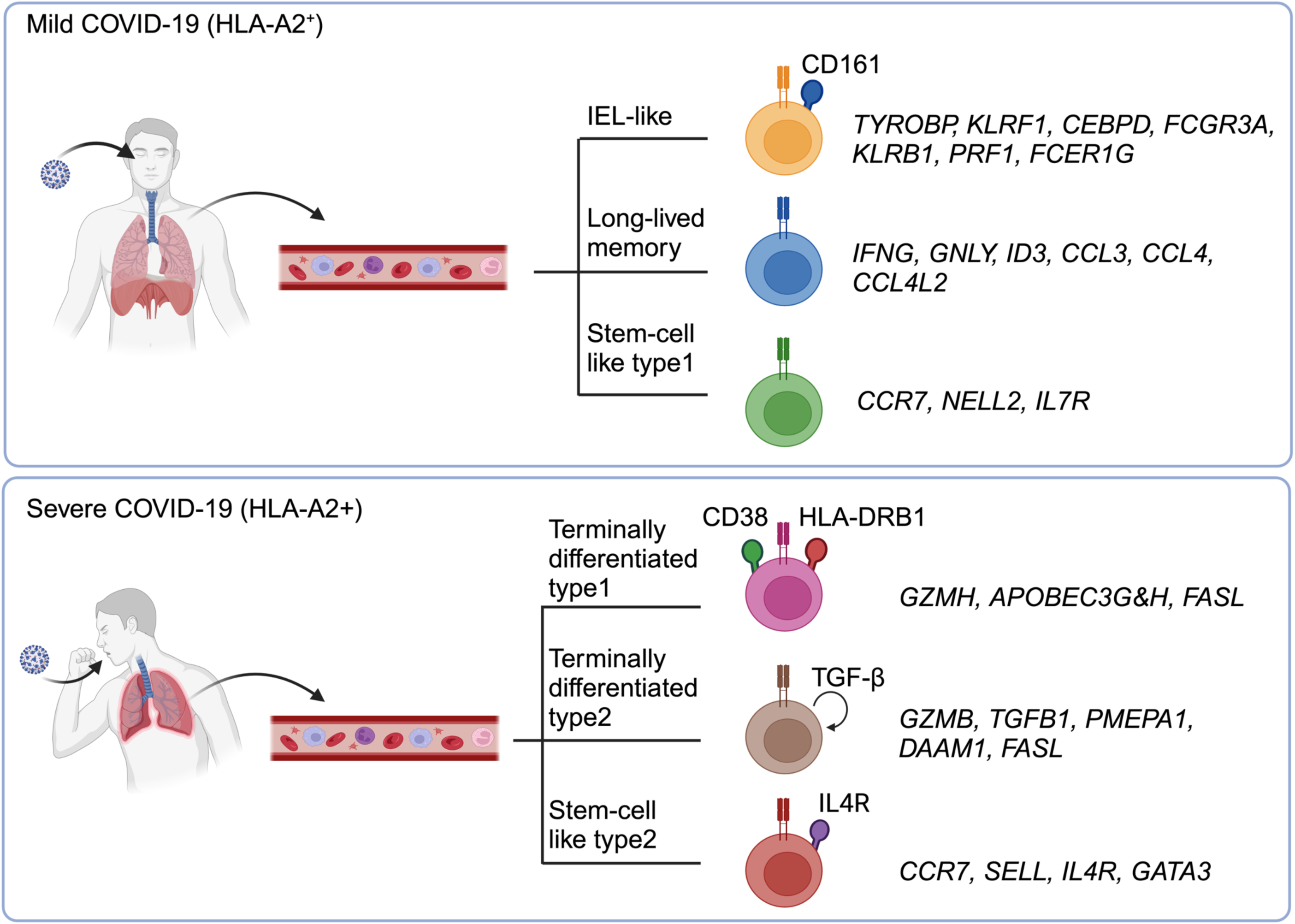

## Introduction

The COVID-19 pandemic caused by *Betacoronavirus pandemicum or* severe acute respiratory syndrome coronavirus 2 (SARS-CoV-2) virus has significantly impacted human health and economiesglobally.^1^;(https://ictv.global/report/chapter/coronaviridae/coronaviridae/betacoronavrus) Despite the initial successes of the vaccines and therapeutics, which have been developed swiftly in the early years of the pandemic, the disease remains a major issue with continued deaths, particularly in at-risk populations. (https://data.who.int/dashboards/covid19/cases) Current vaccines may not elicit durable immune responses. The continued emergence of antibody-resistant variants has been an ongoing challenge for vaccines and antibody-based treatments,^2–5^ and small molecule-based modalities have shown varying levels of efficacy.^6–8^ Furthermore, 65 million people worldwide who are convalescent from COVID-19 display post-acute sequelae in a plethora of organ systems for years after infection.^9^ This phenomenon as well as SARS-CoV-2-triggered immunity, particularly in cell-mediated immunity, are yet to be clearly understood, in order to improve preparedness and aid on the designing of evolution-resilient next-generation interventions.

Adaptive immunity plays a crucial role in preventing SARS-CoV-2 persistence in the upper respiratory tract.^10^ Although most COVID-19 convalescents and vaccinees demonstrate robust antibody responses and T cell immunity,^11^ SARS-CoV-2 is constantly evolving to escape neutralizing antibodies. In this regard, T cells recognize diverse 9-15 linear peptides from SARS-CoV-2, with thousands of unique CD4^+^ and CD8^+^ T-cell epitopes identified to-date.^12–29^ These epitopes are found within virtually all proteins from the virus.^12–28,30^ This broad recognition, coupled with population diversity in HLA polymorphism, makes unlikely evasion of the T-cell response through epitope mutation scarce.^16,30^ Importantly, recent studies^31,32^ indicate that T-cell responses alone may sufficiently control COVID-19 severity as exemplified by evaluation of B cell-deficient patients or animal models. Disease severity correlates with decreased or dysfunctional T-cell levels, including lymphopenia, independent of neutralizing antibody response.^33^ Finally, the inclusion of T-cell epitopes into COVID-19 vaccine designs have been shown to improve their efficacy.^34^

Recent findings demonstrate that memory T-cell responses remain highly cross-reactive two years after initial infection.^11^ Long-term memory T cells, including stem cell memory T (T_scm_) and CD45RA re-expressing terminally differentiated (T_emra_) cells, can persist beyond one year.^35^ To better understand memory CD8^+^ T-cell responses and the memory types associated with COVID-19 severity,^24,36–38^ we conducted single-cell transcriptomics, proteomics, and trajectory analyses of circulating HLA-A2-restricted CD8^+^ T cells recognizing six SARS-CoV-2-specific epitopes from RNA-dependent RNA polymerase (RdRp), Helicase and Spike protein. For that purpose, we utilized peripheral blood mononuclear cells (PBMCs) from unvaccinated HLA-A2^+^ patients with mild, moderate, and acute severe COVID-19, as canonical HLA-A2 allele:02:01 is the most common HLA type prevalent worldwide.^39^ Our analyses revealed, for the first time, enriched SARS-CoV-2-specific effector memory CD8^+^ T-cell types, including cytotoxic long-lived memory T cells and innate-like *KLRB1*^+^ (CD161^+^) CD8aa^+^ T cells, predominantly associated with mild COVID-19 cases having the same HLA-class I allele. Single cell trajectory analysis demonstrated distinct T-cell differentiation pathways between mild and severe cases. In severe COVID-19 patients, naïve CD8^+^ T cells against SARS-CoV-2-specific epitopes resulted in premature termination without proliferation, and different late dysfunctional T-cell subtypes were predominantly observed. These findings emphasize the importance of T-cell mediated immunity in SARS-CoV-2 infection within an HLA-A2-restricted context.

## Results

### Features of unvaccinated and acute HLA-A2^+^ COVID-19 patients with mild, moderate or severe disease status

As our goal was to interrogate the relationship between SARS-CoV-2-directed T-cell responses and phenotypes and disease severity, we utilized a cohort of archived acute (within 4 weeks post diagnosis) COVID-19 patient samples from very early in the pandemic, allowing us to avoid the potential of a confounding effect by prior infections and/or vaccinations. Furthermore, as HLA-class I polymorphism affects the outcome of COVID-19,^40^ we chose to specifically restrict our analyses to only patients with the HLA-A2 allele, so that we could directly link the observed responses to disease severity, without HLA-class I allele variation interference. This allele was chosen as it is present in 40-50% of the global population.^39^ Using PCR genotyping and flow cytometry phenotyping, we identified 42 HLA-A2^+^ samples, which were used for our study (**Table S1**). These patients were then classified based on the disease status, mild, moderate or severe, according to criteria consistent with the WHO Clinical Progression Scale (**Figure 1A** and **Table S2**).^41^ The detailed demographic information of the patients includes the pre-existing medical conditions, COVID-19 treatment, and the outcome of severe patients (**Figures 1A and 1B**). Importantly, age was a critical factor that determines disease severity and was associated with various chronic inflammatory diseases (**Figures 1C** and **1D**).

**Figure 1.**
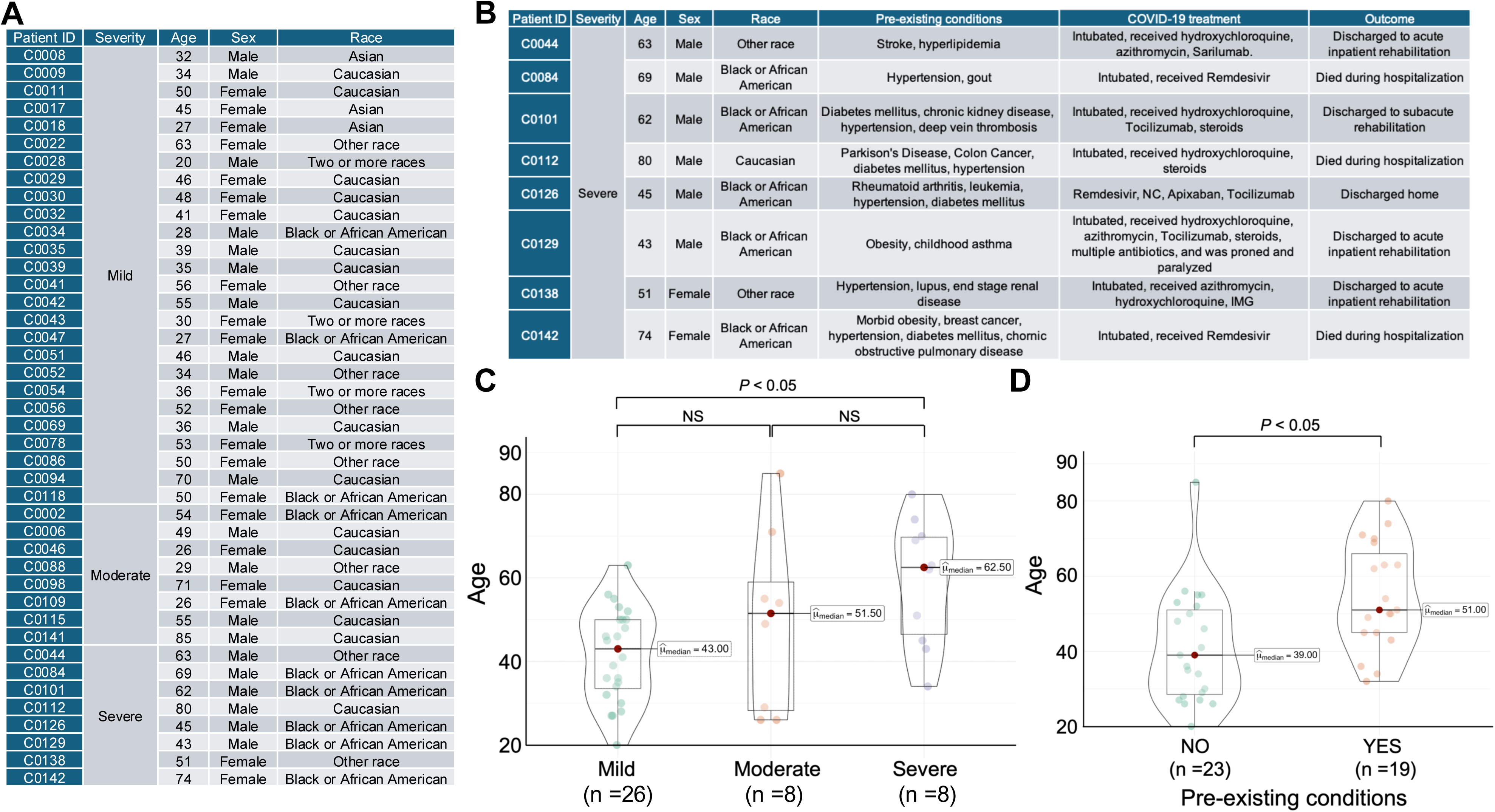
Features of HLA-A2^+^ COVID-19 patients across disease severity. (A) Disease severity and demographic information for 42 HLA-A2^+^ COVID-19 patients. (B) Detailed information of 8 HLA-A2^+^ severe COVID-19 patients (as indicated). (C) Box and violin plots of age in mild, moderate and severe COVID-19 patients and comparison of its median among disease severity. A statistical value (p < 0.05): Mann–Whitney U test. NS: Not Significant. (D) Box and violin plots of age with or without pre-existing conditions and comparison of its median. A statistical value (p < 0.05): Mann–Whitney U test.

### Identification of peptides that encode HLA-A2-restricted SARS-CoV-2-specific CD8^+^ T cell epitopes

To study SARS-CoV-2-specific T-cell responses, we first set out to identify HLA-A2-restricted CD8^+^ T-cell epitopes from SARS-CoV-2 genome. Thirty-one candidate peptides were identified by NetMHCpan-4.0 and chosen based on their HLA-A2-binding score and hydrophobicity (**Figure 2A and Table S3**). These peptides were then used to determine the reactivity of SARS-CoV-2-specific HLA-A2-restricted CD8^+^ T cells derived from HLA-A2^+^ COVID-19 patient PBMCs (**Figure 2A** and **Table S4**). Based on their prevalence score (number of patients among 24 patients who responded to the peptide by ELISpot assay), we selected 11 peptides, which are highlighted in red in **Table S4**, to be used for further studies. Eight peptides were derived from the non-structural proteins (nsps) papain-like protease (PLpro), SARS-unique domain (SUD-M), RNA-dependent RNA polymerase (RdRp), and the 3’ to 5’ exoribonuclease (ExoN), and the other 3 peptides were from the spike protein (**Figures 2B and 2C**). These peptides are highly conserved in SARS-CoV-2, even among the currently circulating variants such as XEC, and some even extend homology to MERS-CoV (**Figure 2B and Table S5)**.

**Figure 2.**
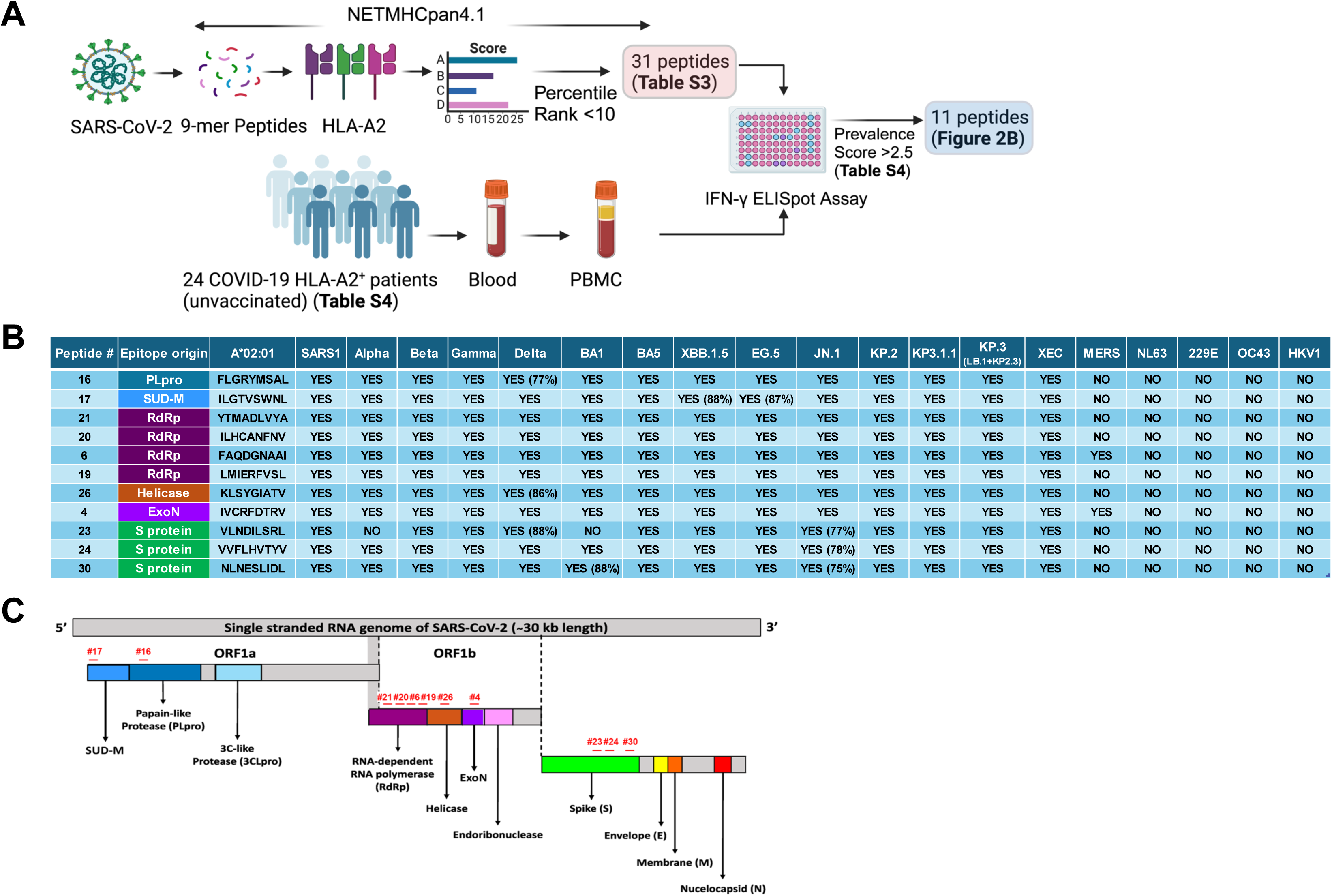
Information of 11 selected HLA-A2 binding 9-mer peptides derived from SARS-CoV-2 genome. **(A)** Workflow of HLA-A2 binding peptide prediction with NETMHCpan4.1 and IFN-γ ELISpot Assay of 24 COVID-19 patient PBMCs with 31 predicted peptides. (B) Homology of 11 selected peptides among SARS-CoV-2 variants and other strains (as indicated). (C) Information of SARS-CoV-2 genome and mapping of 11 selected peptides on its genome.

### SARS-CoV-2-specific CD8^+^ T-cell responses inversely correlate with disease severity in HLA-A2^+^ patients

With our selected peptides, we then conducted IFN-γ ELISpot assays with the entirety of our cohort to assess T-cell responses across 42 HLA-A2^+^ samples of patients with 26 mild, 8 moderate and 8 severe disease states (**Figure 3A**). We observed that these peptides effectively stimulated and elicited CD8^+^ T-cell responses in PBMC samples from patients with mild COVID-19, whereas those with moderate COVID-19 likely had weaker responses. In severe COVID-19 cases, only minimal responses were observed (**Figure 3A**). Nearly all mild cases responded to at least one peptide (20/26; 76.9%). The strongest responses were observed in a mild COVID-19 case (C86) with robust signal against all 11 selected peptides, while PBMCs from other patients responded to varying numbers of peptides. An integrated analysis of the T-cell responses within each of the disease statuses showed a statistically higher response in the mild cohort than the severe (*P* < 0.01) (**Figure 3B**). Mild cases trended higher than moderate cases, and moderate cases trended higher than severe cases, but did not reach a statistical significance. For a subset of the patients for which samples were available for multiple timepoints, we conducted a longitudinal analysis of the CD8^+^ T-cell response. In contrast to neutralizing SARS-CoV-2 antibodies which have been shown to rapidly decay after elicitation,^42,43^ we observed durable responses against all the peptides over the timeframe examined (∼4 months post-diagnosis), as described elsewhere.^44^ In one case (C51), responses continued to increase even up to 104 days post-diagnosis, before beginning to wane (**Figure 3C**). Collectively, these results demonstrate that CD8^+^ T-cell responses broadly target SARS-CoV-2 proteins, are inversely correlated to disease severity, and are sustained for months, affirming their protective function.

**Figure 3.**
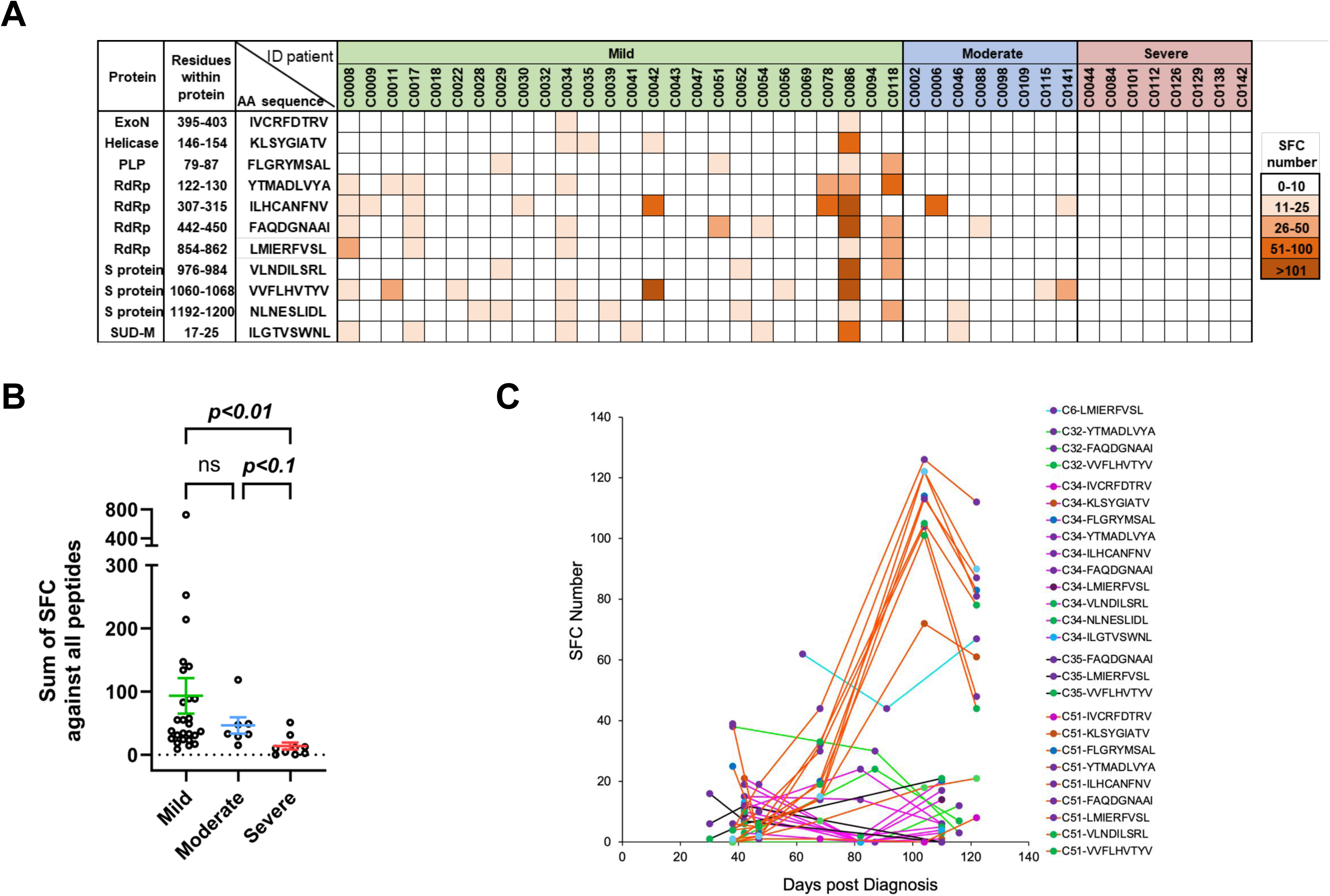
IFN-γ ELISpot of CD8^+^ T cells in HLA-A2^+^ COVID-19 patient PBMCs with selected SARS-CoV-2 peptides. (A) Cross-sectional ELISpot assay of HLA-A2 restricted CD8^+^ T cell responses against 11 selected peptides in 26 mild, 8 moderate and 8 severe COVID-19 patient PBMCs, respectively. (B) Dot plots of a total sum of SFC against a total of 11 selected peptides in mild, moderate and severe COVID-19 patients, respectively and comparison of its median among disease severity. A statistical value (p < 0.01): Mann–Whitney U test. ns: not significant. (C) Longitudinal ELISpot assay of HLA-A2 restricted CD8^+^ T cell responses against selected peptides in mild and moderate COVID-19 patient PBMCs (as indicated).

As a significant component of SARS-CoV-2 adaptive immunity is dictated by antibodies, we next set out to interrogate their role in COVID-19 severity. We conducted ELISAs against the nucleoprotein (NP) and spike, using matched serum samples for 42 patients in our cohort. These antigens were chosen as they have been shown to be the major immunodominant targets for SARS-CoV-2 antibodies, with spike-directed antibodies being a majority of the neutralizing response.^43,45–47^ Across both antigens, strong serum titers tended to be associated with moderate and severe patients, while relatively weak serum titers were observed in mild patients (**Figure S1A**). For the spike, the EC50 in severe patients was significantly higher than in mild patients (*P* < 0.0001) (**Figure S1B**), consistent with a prior study.^48^ We then examined the relationship between CD8^+^ T-cell and humoral responses in disease severity (**Figure S1C**). Although we did not find any correlation between cellular-mediated and humoral immunity, strong cellular immune responses seemed inversely correlated with disease severity regardless of the degree of antibody titer (**Figure S1C**).

### Characteristics of SARS-CoV-2 peptide-loaded tetramer and pentamer positive HLA-A2-restricted single CD8^+^ T cells

Having confirmed SARS-CoV-2 specific peptide reactions with PBMCs from mild, moderate or severe COVID-19 patients, respectively, we next characterized single HLA-A2-restricted CD8^+^ T-cell types that specifically recognize SARS-CoV-2-specific epitopes using selected SARS-CoV-2 peptide-loaded tetramer (Tet) and pentamer (Pent) cocktail that comprised 6 peptides, R_307-315_ (ILHCANFNV), R_442-450_ (FAQDGNAAI), R_854-862_ (LMIERFVSL), S_976-984_ (VLNDILSRL), S_1192-1200_ (NLNESLIDL), H_146-154_ (KLSYGIATV), all of which responded to 11 patient PBMCs (**Figure 4A** and **Figure S2A**). Tet^+^ or Pent^+^ live HLA-A2-restricted CD8^+^ T cells were sorted and subject to the 10x flow (**Figure 4A**). As a result, the UMAP from Tet^+^ or Pent^+^ HLA-A2-restricted CD8^+^ T cells showed diverse T-cell types, in which CITE-seq revealed distinct populations between mild and severe COVID-19 patients (**Figure 4B**). Over 90% of the populations with unique gene signatures, cytotoxic T cells (T_cyto_: *FCGR3A^hi^*, *GNLY^hi^*), effector T cells (T_eff_: *GNLY*, *GZMB*, *PRF1*), trafficking T cells (T_traf_: *CCR6*, *LTB*), developmental T cells (T_dev_: *CD27*, *KLRG1*) and activated T cells (T_act_: *NFKBIA*, *MXRA7*) were observed in patients with mild COVID-19 (**Figures 4B-4D**). A majority of T_cyto_, T_eff_, T_act_ and T_dev_ cell subtypes were from specific patients, respectively (C86: 90% of T_cyto_; C86: 96% of T_eff_; C78: 69% of T_act_; C86: 87% of T_dev_), indicating that those cell types were unique among patients with mild COVID-19 (**Figure 4E**). Although more than 50% of single cells in the cell types with innate-like gene signature (T_in-like_: *FCER1G*, *PRF1*, *GNLY*), stem cell-like memory signature (T_scm1_: *LTB*, *CD27*, *CCR7*) or resident memory signature (T_rm_: *ITGA1*, *IL7R*) were from mild COVID-19 patients (**Figure 4D**), the cell composition of each patient within these clusters was not predominant (<50%) (**Figure 4E**), suggesting that those cell types were likely to be commonly observed among patients exhibiting mild COVID-19 symptoms (**Figures 4D and 4E**). Interestingly, early antigen-experienced T-cell subtype (T_scm2_: *IL4R*, *CCR7, SAMSN1, IL7R, LTB*) was enriched in patients with severe disease, particularly (C101) (**Figures 4B and 4C and 4E**), suggesting that such early antigen-experienced T cells may have culminated into dysfunctional cells. In contrast, the naïve T-cell type (T_naive_: CCR7^hi^) with high levels of CD45RA expression was dominated by neither mild nor severe COVID-19 patients (**Figure 4D** and **Figure S2D**). Those results indicate that the T_naïve_ cell type could have two distinct lineages depending on patient conditions. The recruiting T-cell type (T_rec_: *CCL4L2*) and terminally differentiated T-cell subtypes (T_td1_ and T_td2_) were preferentially observed in severe COVID19 patients (**Figure 4D**). Over 95% cell populations, two different terminally differentiated T-cell (T_td_) subtypes with unique gene features (T_td1_: *EOMES*, *HLA-DRB1*; T_td2_: *SAMSN1*, *ZNF683*) were specifically observed in the patients C101 and C138 with severe COVID-19, respectively (**Figure 4F**). The CITE-seq analysis revealed that almost whole single T cells were composed of CD69^+^ cells (>99%), CD137^+^ cells (>95%), and CD69^+^CD137^+^ cells (>95%), known as antigen-specific markers, validating that all those CD8^+^ T cells were likely to be SARS-CoV-2 epitope-specific cells as previously described.^29^ Effector memory T-cell types, T_cyto_ and T_eff_ possessed the cells with most prominent expression of CD45RA, indicative of terminally differentiated CD45RA^hi^ (T_emra_) cells (**Figure S2D**), suggesting that those cell types are likely to have a lineage distinctly from T_td_ cell subtypes.

**Figure 4.**
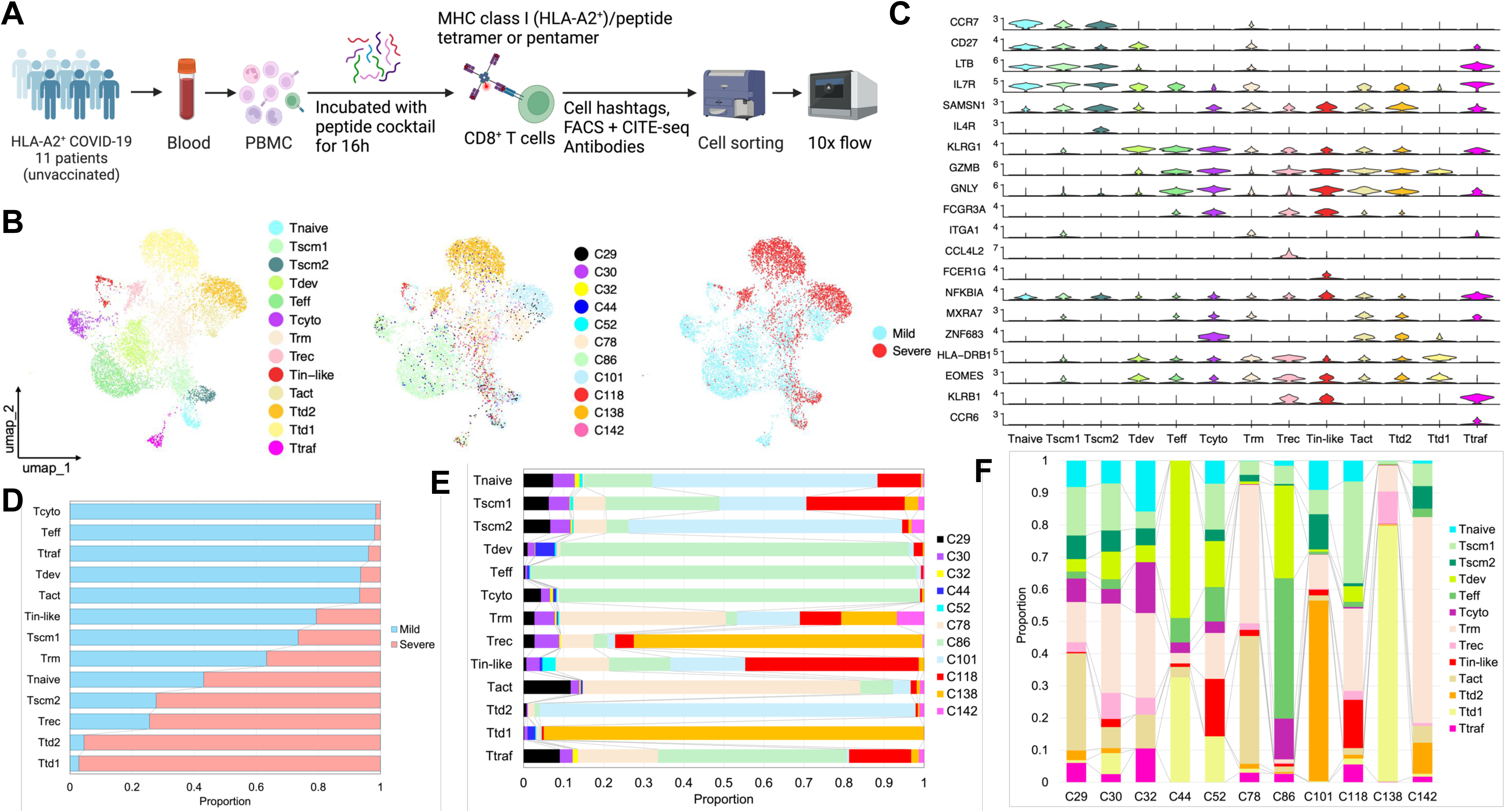
Overview of SARS-CoV-2 derived epitope-specific A2^+^ CD8^+^ single T cell types. (A) Workflow of the single-cell experiment. (B) UMAP plots of the identified CD8^+^ T cell clusters (left), each patient distribution (middle), and distinct populations from mild and severe COVID-19 patients (right). (C) Violin plots of the selected genes (as indicated) in each identified CD8**^+^** T cell cluster. (D) Proportion of single cells from mild and severe COVID-19 patients in each cluster. (E) Distribution of single cells from all patients in each cluster. (F) Distribution of the whole single cells in each patient.

### Immunodominant effector CD8^+^ T-cell types associated with mild COVID-19 status

Of the identified effector T-cell types, we found potent effector SARS-CoV-2 epitope-specific HLA-A2-restricted CD8^+^ T-cell types (T_in-like_, T_rec_, and T_traf_) with unique gene features that predominantly contained *KLRB1*^+^ (CD161^+^) cells (**Figure 5A and Figure S3A**). The T_in-like_ cluster possessed many CD161-expressing CD8^+^ T cells that lacked CD8B expression (**Figures S3B and S3C**), indicating that those T-cells shared the features of CD8αα intraepithelial T lymphocytes (IELs), therefore renamed the T_IEL-like_ cluster. This cluster included less exhausted cytotoxic cells shared with the genes that natural IELs and NK cells expressed (*CEBPD*, *CLIC3*, *FCER1G*, *FCGR3A*, *IL2RB*, *KLRB1*, *KLRC1*, *KLRC2*, *KLRC3*, *KLRD1*, *SAMD9L*, *SH2D1B*, *TYROBP*) (**Figures 5A and 5B**).^49^ Notably, this T-cell type was enriched in mild but not severe COVID-19 patients (**Figure 5C**). The T_traf_ cluster consisted of the cells with a higher expression of CD161 than the other clusters (T_IEL-like_, T_rec_), most of which expressed MAIT-associated genes (*LTB*, *CEBPD*) and Th17-related genes (*CCR6*, *CXCR6*, *RORA*), suggesting that this cluster shared with MAIT gene signature, so called the T_MAIT-like_ cluster (**Figure 5A and Figure S3A**).^50^ Thus, we found unique cell types with the features of IELs and MAIT that likely reside within the epithelial layer of mucosal and barrier tissues and cytotoxic *KLRB1*^+^ (CD161^+^) CD8αα CD8^+^ T cells could be highly potent for efficiently eliminating SARS-CoV-2 virus. Interestingly, although the T_rec_ cluster mostly consisted of populations from severe COVID-19 patients that expressed unique representative genes related to T-cell recruitment such as *CCL3L1*, *CCL4L2*, and *CCL4* (**Figures 4D and 5D**),^51^ we also found a minor but potent cytotoxic population in this cluster that primarily expressed the *IFNG* gene (**Figure 5E**). To enrich this population, we conducted unsupervised sub-clustering within the T_rec_ cluster. As a result, this cluster was subdivided into 2 distinct clusters (C1, C2) where we confirmed a high density of *IFNG* expression in the C1 cluster, while the C2 cluster mainly contained *KLRB1*^+^ (CD161^+^) cells (**Figure S3D**), suggesting that the C1 cluster contained a unique population independently of *KLRB1*^+^ (CD161^+^) cells. Interestingly, the C1 cluster contained the cells that expressed the genes *LTB* and *ID3* involving long-lived memory T-cells with effectors including *IFNG* (**Figure 5F and Figure S3D**),^52^ while the C2 cluster likely primarily had the cells with high levels of CD45RA and the genes (*EOMES*, *FGFBP2*), indicative of enriched T_emra_ cells (**Figures S3E and S3F**).^53^ Notably, compared to the C2 cluster (>90% cells from severe COVID-19), the C1 cluster mainly accounted for populations from diverse patients with mild COVID-19 status (>74%) (**Figures 5G and 5H**) and long-lived memory CD8^+^ T-cell population in the T_rec_ cluster was significantly enriched in mild COVID-19 patients (**Figure 5I**). Taken together, the C1 cluster possessed potent cytotoxic long-lived memory HLA-A2-restricted CD8^+^ T-cells that potentially recruited other effector T-cells and were therefore effectively able to kill SARS-CoV-2 virus with cytotoxic activity as well as effector *KLRB1*^+^ (CD161^+^) CD8^+^ T cells.

**Figure 5.**
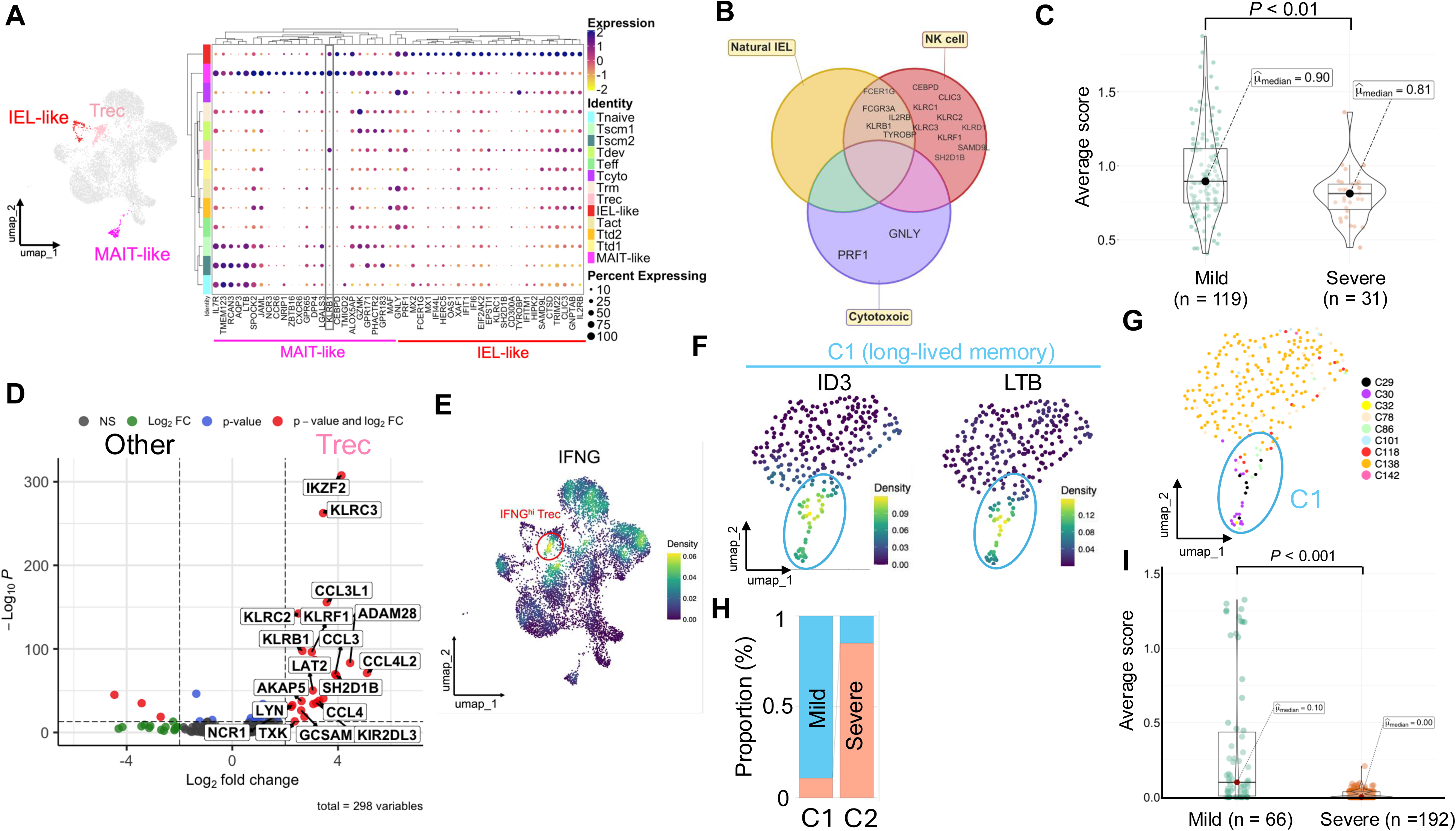
Transcriptomic profiling of unique potent cytotoxic SARS-CoV-2 derived epitope-specific A2^+^ CD8^+^ T cells. (A) UMAP plots highlighted in the T_IEL-like_, T_rec_, and T_MAIT-like_ clusters (left) and gene dot plots in each cluster (as indicated) (right). (B) Venn diagram of natural IEL, NK cell and cytotoxic gene markers (as indicated). (C) Comparison of box and violin plots of average score (mean of top 30 genes in the T_IEL-like_ cluster in **Table S8**) with median points and a statistical value (p < 0.01): Mann–Whitney U test. (D) Volcano plots of differentially expressed genes within the T_rec_ cluster. 18 top genes (as indicated) were highlighted. (E) Density plots of *IFNG* gene expression. The population with a higher density was circled in red (F) Density plots of *ID3* and *LTB* gene expression within the T_rec_ cluster highlighted at the C1 cluster (circled in blue). (G) UMAP plots of patient distribution within the T_rec_ cluster highlighted at the C1 cluster (circled in blue). (H) Proportion of single cells from mild and severe patients in the C1 and C2 cluster within the T_rec_ cluster. (I) Comparison of box and violin plots of average score (mean of top 30 genes in the C1 cluster in **Table S8**) with median points and a statistical value (p < 0.001): Mann–Whitney U test.

### Terminally differentiated effector CD8^+^ T-cell subtypes associated with severe COVID-19 status

Having confirmed highly potent cytotoxic gene signature of HLA-A2-restricted CD8^+^ T-cell types, we next explored antigen-experienced HLA-A2-restricted CD8^+^ T-cell types with unique gene signature and cell surface expression that may no longer function as a cytotoxic T-cell type or be in transition into terminally differentiated cells including exhausted-like cells. We observed highly activated and possibly terminally differentiated two different T-cell subtypes (T_td1_, T_td2_) with original gene features (**Figure 6A**). Interestingly, both cell subtypes expressed FAS Ligand with high levels of effector molecules (**Figure 6B**), suggesting a terminal activation was likely to be elicited in those T-cells. In consistent with a prior study,^54^ one of the terminally differentiated T-cell subsets, T_td1_ included the cells with high levels of CD38 and HLA-DR expressions (CD38^hi^ HLA-DR^hi^) with severe COVID-19 (**Figure S4A**). Nonetheless, the T_td1_ cell subtype showed a previously uncharacterized gene signature with gene markers of cytotoxicity (*GZMB*, *GZMH*) and T-cell infiltration (*APOBEC3H* and *APOBEC3G*) (**Figure 6A and Figures S4B-C**).^55^ This suggests that tissue-infiltrating CD38^hi^ HLA-DR^hi^ effector CD8^+^ T-cells in this cluster were likely to have become late dysfunctional since these HLA-A2 restricted CD8^+^ T cells encountered SARS-CoV-2 specific peptides under a highly inflammatory condition. Interestingly, the other terminally differentiated T_td2_ cell subtype showed a relatively higher level of TGF-β expression (**Figure 6C**). Moreover, this T-cell subtype upregulated the TGF-β signaling-related genes *PMEPA1* and *DAAM* that potentially induced ROS involved in cell damage,^56,57^ and the T-cell exhaustion related gene *ZNF683* and protein PD-1 (**Figure 6E**). These results indicate that TGF-β signaling could suppress its effector T-cell function. In summary, those distinct T-cell subtypes were likely dysfunctional and primarily observed in severe COVID-19 patients (**Figure 6F**).

**Figure 6.**
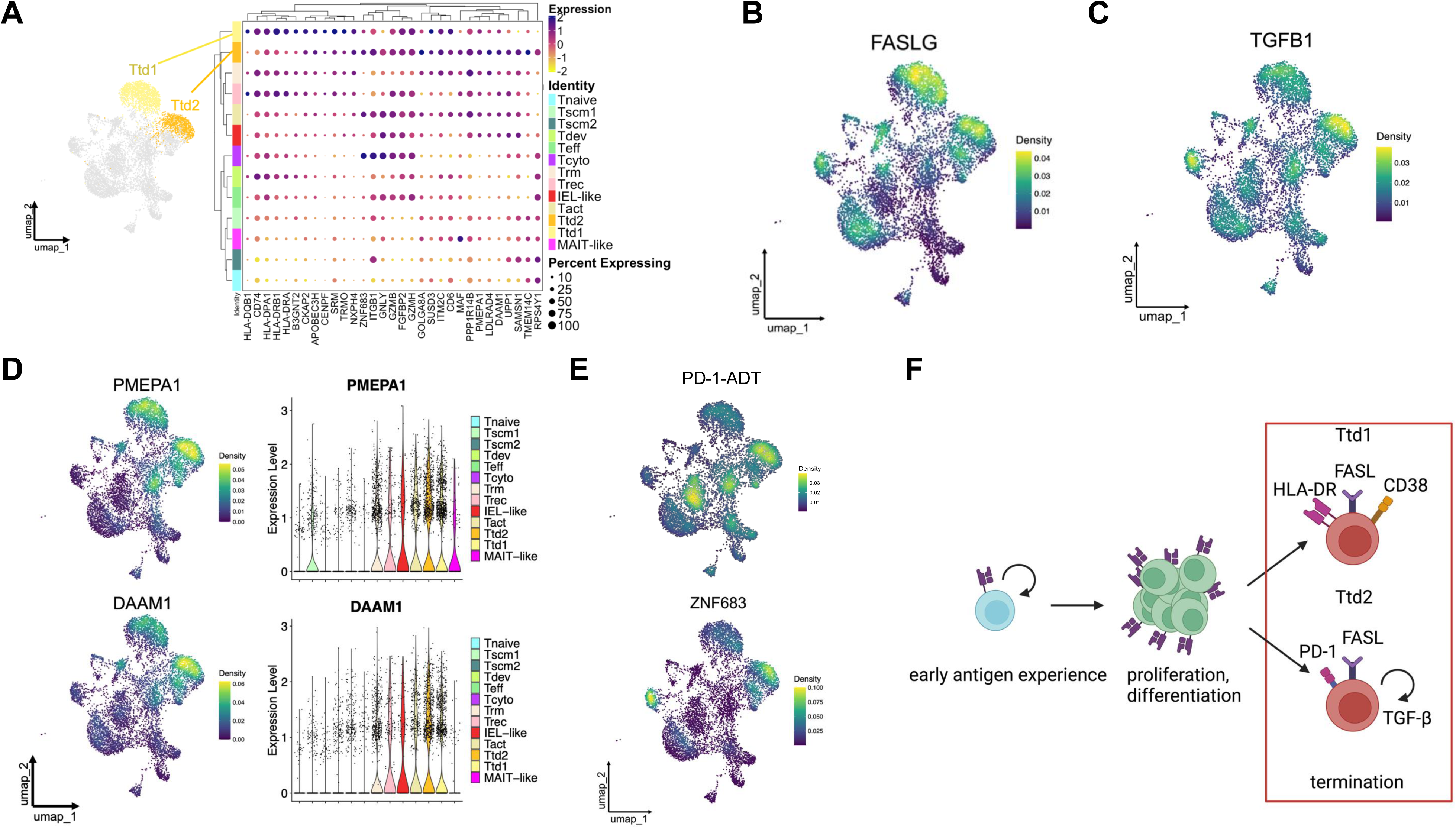
Distinct dysfunctional SARS-CoV-2 epitope-specific A2^+^ CD8^+^ T-cell subtypes. (A) UMAP plots highlighting T cell subtypes (T_td1_, T_td2_) (left) and gene dot plots of the selected genes (as indicated) in each cluster (right). (B) Density plots of *FASLG* gene expression in the UMAP. (C) Density plots of *TGFB1* gene expression. (D) Density plots of *PMEPA1* and *DAAM1* gene expression in the UMAP, respectively (left). Violin plots of single cells with *PMEPA1* and *DAAM1* gene expression, respectively (right). (E) Density plots of PD-1-ADT and *ZNF683* gene expression in the UMAP. (F) The flow of possible CD8^+^ T cell differentiation pathways into two highly activated cell subtypes in the UMAP.

### Distinct SARS -CoV-2-specific stem cell-like CD8^+^ T-cell subtypes associated with COVID-19 severity

We next investigated the fate of HLA-A2-restricted naïve CD8^+^ T-cells against SARS-CoV-2 epitopes. Notably, we found distinct SARS-CoV-2 epitope-specific stem-like HLA-A2-restricted CD8^+^ T-cell subtypes (T_scm1_, T_scm2_), while one naïve T-cell type was only observed in the UMAP containing populations from all the patients except the patient C44 (**Figure 4E** and **Figure 7A**). Unsupervised clustering confirmed a relatively higher level of *CCR7* expression as a naïve T-cell marker within the T_naive_ cluster and subsequently its expression was associated with the T_scm1_ and T_scm2_ clusters (**Figure 7B**). Both T_scm1_ and T_scm2_ clusters possessed the cells with expression of naïve cell markers including *SELL*, *LEF1*, and *NELL2*, while T_scm2_ cluster contained the cells with lower levels of expression of the *CD27* gene, known as a proliferation marker (**Figure 7C** and **Figure S5**). These results suggest that upon stimulation, naïve T-cells can be differentiated into 2 different types of stem-like CD8^+^ T cells. Interestingly, T_scm1_ subtype was enriched in the patients with mild COVID-19, while T_scm2_ subtype was predominantly seen in the patients with severe COVID-19 (**Figure 4B**). Furthermore, the T_scm2_ cluster possessed possibly dysfunctional or early terminated cells with its gene signature (*IL4R*, *GATA3*) (**Figure 7D**). Next, to investigate a lineage of those stem-like CD8^+^ T cells, RNA trajectory analysis was performed based on gene expression in the identified T-cell clusters. As a result, in severe COVID-19, naïve CD8^+^ T cells could early terminate into dysfunctional cells as seen in the T_scm2_ cluster (**Figure 7E**). In contrast, in mild COVID-19, naïve CD8^+^ T cells were potentially able to differentiate into various kinds of effector T-cells including resident memory T cells (T_rm_), activated T cells (T_act_), and developmental T cells (T_dev_) via the T_scm1_ cluster. Those T cells were further able to culminate into terminally differentiated T cells such as highly activated, exhausted-like CD8^+^ T and T_emra_ cells including T_cyto_ and T_eff_ (**Figure 7E)**. Indeed, some mild and severe COVID-19 patients exhibited possible distinct differentiation pathways (**Figures 7E and 7F and Figures S6A and S6B**). Thus, the fate of early antigen experienced CD8^+^ T cells could be determined depending on various disease status, i.e. mild versus severe COVID-19 of patients.

**Figure 7.**
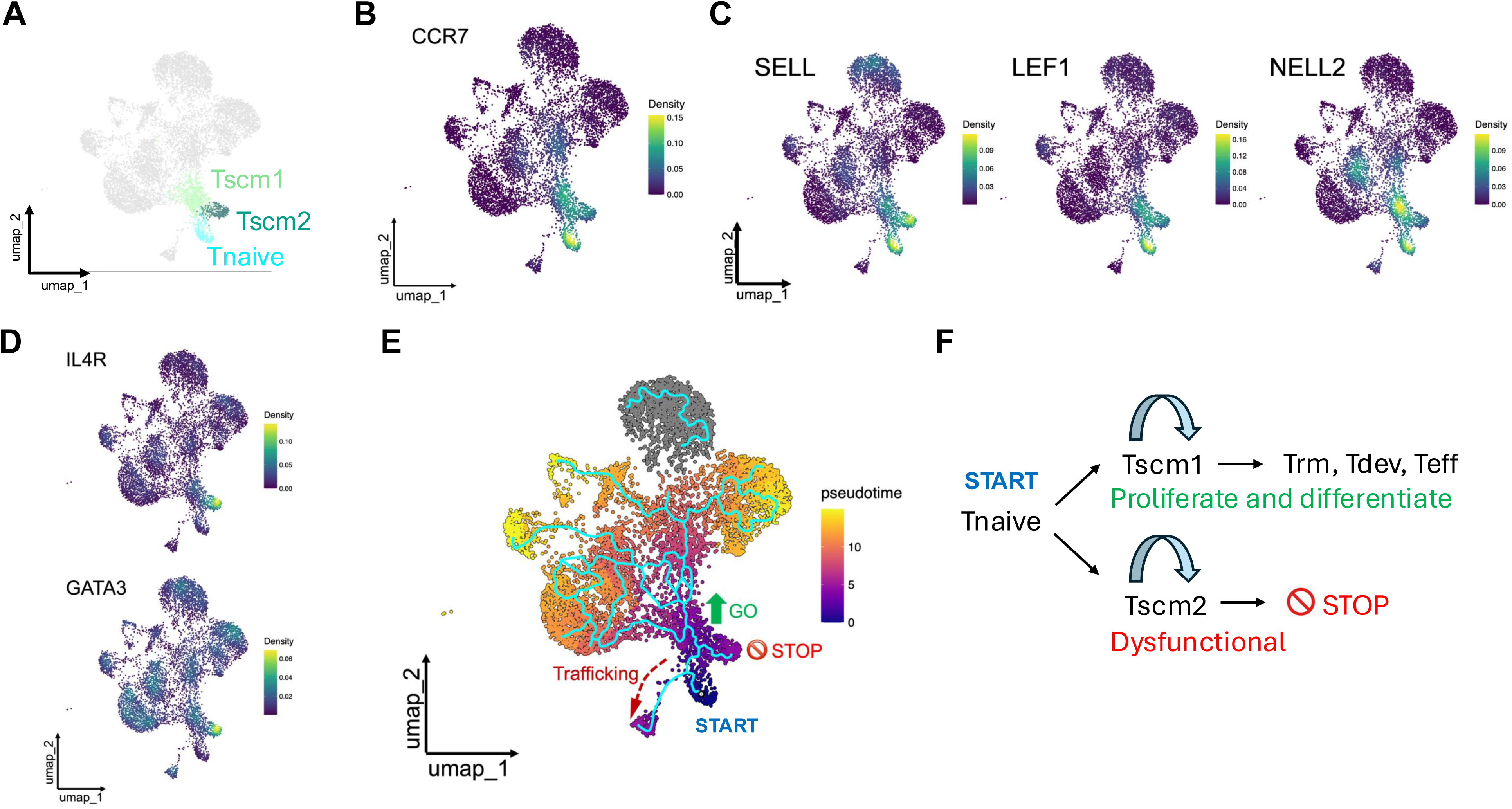
Distinct stem-like and naïve CD8^+^ T-cell types against SARS-CoV-2 epitopes. (A) UMAP plots highlighting T cell types (T_naive_, T_scm1_, and T_scm2_). (B) Density plots of *CCR7* gene expression in the UMAP. (C) Density plots of *SELL*, *LEF1*, and *NELL2* gene expression in the UMAP. (D) Density plots of *IL4R* and *GATA3* gene expression in the UMAP. (E) Single-cell trajectory and pseudotime heat map in the UMAP. “Pseudotime” infers cell differentiation status from naïve to effector T cell types. (F) Possible differentiation pathways of the naïve T cell type.

## Discussion

We observed significantly reduced SARS-CoV-2-specific, HLA-A2-restricted CD8^+^ T-cell responses in severe COVID-19 patients, regardless of their humoral response levels, while these CD8^+^ T-cell responses in mild and moderate patients persisted for up to 4 months post-diagnosis, indicating the significant impact of long-lived SARS-CoV-2-specific memory CD8^+^ T-cell responses on the disease status. Furthermore, we determined that most, if not all, of the immunodominant HLA-A2-restricted CD8^+^ T-cell epitopes are highly conserved across numerous SARS-CoV-2 variants, including the currently circulating XEC variant. This conservation likely occurs because CD8^+^ T-cell epitopes within SARS-CoV-2 genome, unlike B-cell epitopes, are not exposed to immune pressure.^13^ While several effector memory T-cell subsets have been identified, the increasing number of vaccinated individuals complicates efforts to track antigen-experienced memory CD8^+^ T cells generated upon initial SARS-CoV-2 exposure.^58,59^ In this study, HLA-A2^+^ blood samples in unvaccinated patients enabled to delineate the significant importance of T-cell mediated immunity and previously uncharacterized CD8^+^ T-cell types against SARS-CoV-2-specific epitopes and their potentially cytotoxic lineage associated with mild COVID-19 symptoms versus severe disease outcomes.

The impaired T-cell response in severe COVID-19 patients appears to result from both reduced T-cell numbers and the presence of dysregulated or dysfunctional T cells. For instance, dysfunctional spike protein-specific T cells in severe COVID-19 and certain bystander T cells such as lung-homing CXCR4^+^ CD8^+^ T cells was associated with fetal COVID-19 outcomes.^24^ Distinct CD8^+^ T-cell populations against SARS-CoV-2 virus in relation to COVID-19 severity, however, have not been characterized at single-cell resolution.

Our study showed that CD8^+^ T-cell populations, such as T_IEL-like_, T_MAIT-like_, T_dev_, T_cyto_, T_eff_, T_act_, and T_scm1_ were preferentially observed in mild COVID-19 patients. Of these cell types, the T_MAIT-like_ and T_IEL-like_ populations mainly contained *KLRB1*^+^ (CD161^+^) cells that were apparently unique distinctly from the T_dev_ and T_act_ types with more exhausted-like PD-1^hi^ cells and the T_eff_ and T_cyto_ types with CD45RA-reexpressing terminally differentiated, T_emra_ cells and early antigen experienced cells within the T_scm1_. CD161 (*KLRB1*) is human homolog of NK1.1 that is expressed on NK cells, T-cell subsets and NKT cells.^60^ CD161^+^ CD8^+^ cells contain innate-like T cells that are involved in anti-viral responses.^61^ CD8^+^ T cells with a relatively higher expression of CD161 are particularly categorized into MAIT cells.^50^ A prior study indicated that MAIT cells as innate-like T cells played an essential role in protection against SARS-CoV-2 virus, although altered or decreased MAIT cells correlated with disease severity.^62^ Our identified HLA-A2-restricted MAIT-like cell phenotype against SARS-CoV-2-specific epitopes exhibited proliferative markers such as *IL7R* and *LTB* rather than effector functions such as *GZMB*. This cell-type also shared chemokine *CCR6* and *CXCR6* related to Th17-cell signature. Thus, these MAIT-like cells are likely involved in early antigen responses against SARS-CoV-2 virus. In contrast, the T_IEL-like_ type with less CD161 expression contained more cytotoxic gene signature such as *GNLY* and *PRF1* was frequently observed in a mild patient, C118. Interestingly, this cell-type accounted for a majority of CD8A^+^ cells with loss of CD8B expression, a hallmark of intraepithelial T lymphocytes (IELs).^49^ CD8aa^+^ CD161^+^ T cells are also reported to exert as a CD8 memory cell-type.^63^ Hence, our identified T_IEL-like_ cells may not be exactly IELs, as they normally reside in the epithelium of various organs that include lung and upper respiratory tract.^64^ Nonetheless, this cytotoxic innate-like CD8^+^ T-cell type likely plays an essential role in protecting and controlling SARS-CoV-2 infection, given recent studies have presented compelling evidence of the importance of upper-airway memory T cells or lung-resident T cells in the context of SARS-CoV-2 infection in humans.^59,65^ In conclusion, the expansion of HLA-A2-restricted innate-like CD8^+^ T-cell populations in PBMCs could be related to the disease outcome.

In severe COVID-19 patients, we found unique effector T-cell subtypes (T_td1_ and T_td2_) primarily expressing FAS ligand, known as a highly activated T-cell marker.^66^ Such phenotypically distinct T-cell subtypes were enriched in certain COVID-19 patients (T_td1_: C138; T_td2_: C101), suggesting that these T-cell subtypes may have been controlled under the state of chronic inflammatory conditions such as diabetes mellitus and chronic kidney disease (C101) and lupus and end stage renal disease (C138) in SARS-CoV-2 infection. The T_td1_ cells express effector molecules, such as Granzyme B and H, but also cell infiltration and tissue-association markers with high levels of HLR-DR and CD38. These T_td1_ cells were primarily observed in a severe COVID-19 patient, C138. Emerging evidence indicated that HLA-DR^hi^ CD38^hi^ CD8^+^ T-cell subtype was enriched over a long-time period in severe patients infected with SARS-CoV-2.^67,68^ Thus, T_td1_ cells were likely to be highly activated and dysfunctional under tissue-specific inflammatory environments. Interestingly, another T-cell subtype, T_td2_ cells express effector molecules, such as Granzyme B and H and perforin with high levels of TGF-β expression. The effects of TGF-β on T-cell activation vary depending on the state of T cells. Some groups reported that TGF-β promotes effector Th17 differentiation and Treg development in an early stage,^69^ while others concluded that TGF-β signaling inhibits cytotoxic T-cell functions.^70^ T_td2_ cells we identified exhibited activation of TGF-β signaling followed by its downstream upregulation of *PMEPA1* and *DAAM*, indicative of autocrine TGF-β stimulation. Since the gene *PMEPA1* is induced under devastating conditions related to hypoxic stress responses and pro-fibrotic responses and the gene *DAAM* expression involves stress fiber formation that could limit T-cell activation,^71^ T_td2_ cells might have been exhausted or burning out. In fact, T_td2_ cells also express high levels of T-cell exhaustion markers such as PD-1 and ZNF683.

Lineage tracking through transcriptional changes enabled us to trace T-cell fates as they developed into either dysfunctional or potent cytotoxic cells. Notably, we identified an early dysfunctional T-cell subtype, T_scm2_. Our trajectory analysis revealed premature termination of early antigen-experienced CD8^+^ T cells within the T_scm2_ cluster. T_scm2_ subtype expressed both IL-4 receptor and GATA3. While IL-4 receptor seems to play an important role in early CD8^+^ T cell development, ^72^ GATA3 is known to promote CD8^+^ T cell dysfunction in tumor immunity.^73^ Consequently, the T_scm2_ cells likely failed to proliferate and differentiate into effector cells upon SARS-CoV-2 antigen recognition. Although the mechanisms underlying the enrichment of these dysfunctional CD8^+^ T-cell subtypes in severe COVID-19 patients remain unclear, it is conceivable that advanced age and/or pre-existing medical conditions in severe patients may have compromised CD8^+^ T-cell functionality as described somewhere.^74^

### Limitations of the study

While our ELISpot assays have determined the number of CD8^+^ T cells that secrete IFN-γ in response to stimulation by SARS-CoV-2 peptides, our single cell analyses detected prominent IFN-γ expression only in part of antigen-experienced effector CD8^+^ T cells. This apparent discrepancy likely stems from differences in assay timing post peptide stimulation. The IFN-γ spots in ELISpot assays represent cumulative cytokine production over a 24-hour stimulation period. In contrast, for single cell analyses, we could only conduct measurements after 16 hours post-stimulation with the peptide cocktail, as the TCR-CD3 complex undergoes downregulation between 2-12 hours after activation.^75^ During this period reduced TCR expression prevents reliable tetramers/pentamers detection of antigen-specific T cells.

For our single-cell analysis, we could not test all 42 patients due to limited PBMC sample availability. The ELISpot assay requires a minimum of 2 x 10^7^ PBMCs to run in duplicate. Only 1 patients (7 mild and 4 severe patients) had sufficient samples with more than 3-4 vials each containing 1 x 10^7^ PBMCs per vial. These samples permitted us to perform the tetramer/pentamer assays, followed by the T-cell sorting and single-cell analysis.

## Methods

### Ethics statement

This protocol was approved by the Columbia University Irving Medical Center IRB (#AAAS9722)

### Sample collection

Frozen peripheral blood mononuclear cells (PBMCs) from the acute and convalescent phase of SARS-CoV2 infection were obtained from the COVID-19 Persistence and Immunology Cohort (C-PIC) study led by investigators in the Division of Infectious Diseases, Department of Medicine at Columbia University Irving Medical Center. Patients were diagnosed either upon admission to the hospital or during outpatient visits. Three to five cryovials, each containing 10^7^ PBMCs, from each COVID-19 patient were collected at multiple timepoints within the first year after infection, processed and frozen. For acute patients, PBMCs were collected between 1 week and 4 weeks after diagnosis, and for our longitudinal study, PBMCs were collected up to 120 days post diagnosis. Before thawing the PBMCs, we extracted a small piece of the frozen cell pellet using a sterile disposable Corning V-scoop/spoon. The piece was dissolved in a sterile tube containing RPMI medium, followed by the addition of Enhancer Solution and Proteinase K. After vortexing the mixture, DNA was extracted using DNA binding beads, following the manufacturer’s instructions (MagMAX™ DNA Multi-Sample Ultra 2.0 Kit, Thermo Fisher Scientific). We then identified HLA-A2^+^ patients using two methods: 1) Genotyping: Twenty-seven patients were identified as HLA-A2^+^ through this method. 2) Phenotyping: We incubated one million PBMCs with PE-labeled anti-HLA-A2 antibody (BioLegend), followed by flow cytometric analysis using BD LSRII. Fifteen patients were identified as HLA-A2^+^ using this approach. In total, we identified 42 HLA-A2^+^ individuals out of 77 COVID-19 patients, as shown in **Table S1**. The percentage of HLA-A2^+^ patients was 54.5%, which aligns with the previously mentioned global prevalence of HLA-A2. The grade of the disease severity of the 42 HLA-A2^+^ COVID-19 patients was determined based on the guidelines listed in **Table S2**.

### HLA typing

We determined the HLA-A2 allele of each PBMC sample by genotyping or phenotyping. For genotyping, we cut small pieces (10 µL) of cell pellet from a vial that contains frozen human PBMC sample from convalescent COVID-19 patients, using sterile disposable rigid Corning V-scoop/spoon (Millipore Sigma), which was then dissolved into sterile tubes with 40 µL of RPMI, adjust final volume up to 50 µL. In a 96-deep-well plate, we first added 5 µL of Enhancer Solution (Thermo Fisher), followed by adding 50 µL of RPMI containing PBMC sample, and lastly 5 µL of Proteinase K (Thermo Fisher) was added to each well and sealed. The plate was vigorously vortexed for at least 30 s and then incubated for 20 min at 65 °C. DNA was then purified by using DNA-binding beads (Thermo Fisher). Samples were then typed by PCR using sequence-specific oligonucleotide probes for the HLA-A gene (American Red Cross Blood Services). Convalescent COVID-19 patient samples were handled in a biosafety level-3 (BSL-3) lab. For phenotyping, after thawing the PBMCs, we put 1 x 10^6^ cells aside, which were stained with PE-labeled anti-HLA-A2 antibody (BioLegend), followed by flow cytometric assay, as described.^76^

### Peptides

CD8^+^ T-cell epitopes within the SARS-CoV-2 genome were identified by utilizing NetMHCpan-4.1 and then manually curated based on binding score, hydrophobicity, and homology to other coronaviruses.^77^ The chosen peptides were synthesized by Peptide 2.0 Inc., at 85% purity.

### Homology analysis of SARS-CoV-2 9-mer peptides against SARS-CoV-2 variants and other coronaviruses

To analyze the conservation of the selected 9-mer peptides in SARS-CoV-2, SARS-CoV-1, MERS, and other common human coronaviruses, including NL63, 229E, and HKU1, we first identified the position of each protein sequence in a reference SARS-CoV-2 genome, *hCoV-19/Wuhan/WIV04/2019* (Accession ID: EPI_ISL_402124), downloaded from the GISAID database [https://gisaid.org/]. We used Nextclade [https://clades.nextstrain.org/] to identify each protein sequence of interest from viral genome sequences obtained from both GISAID and the BV-BRC website [https://www.bv-brc.org/]. For each clade of SARS-CoV-2 and SARS-CoV-1, we collected 100 sequences from both the earlies and most recent available time points. This approach provided a dataset of 200 sequences per lineage to analyze mutation frequencies over time. For MERS and the other viruses, which have few genome sequences available, we utilized one representative genome per virus. Sequences of each protein were then search for the 9-mer motifs to calculate their occurrence.

### IFN-γ ELISpot assay

IFN-γ levels from patient PBMCs upon stimulation by specific peptides were quantified by ELISpot assay. MultiScreen-IP filter 96-well plates (MAIS4510; Millipore Sigma) were prewashed quickly (<60 sec) with 70% ethanol, followed by been washed the wells with sterile water and coated with anti-human-IFN-γ antibody (MABTECH) overnight at 15 µg/mL in PBS at 4 °C. The following day, wells were washed twice with PBS, three times with DMEM + 10% FCS, and then blocked for 2 h at 37 °C with 200 µL of DMEM + 10% FCS. Wells were then washed once with DMEM + 10% FCS + rIL-2 (10 ng/mL), and then 500,000 PBMCs in 100 µL of DMEM + 10% FCS + rIL-2 (10 ng/mL) and the appropriate peptide(s) at 10 µg/mL in 100 µL of DMEM + 10% FCS + rIL-2 (10 ng/mL) were added to each well. Plates were then incubated for 18 to 22 h at 37 °C under 5% CO_2_. Wells were then washed twice with PBS, four times with PBS + 0.05% Tween-20 (PBST), and then incubated with 100 µL/well of biotinylated anti-human-IFN-γ (MABTECH) at 2 µg/mL in PBST for 2 h at RT. Wells were washed four times with PBST, and then incubated with 100 µL/well of HRP-Avidin (BioLegend) at 1:800 dilution in PBST for 45 min at RT. Wells were washed four times with PBST, twice with PBS, and then spots were developed by adding AEC ELISpot Substrate (BD) for 15 min at RT and quantified by microscopy. Convalescent COVID-19 patient samples were handled in a biosafety level-3 (BSL-3) lab.

### ELISA

The S2P spike trimer protein and nucleocapsid protein (NP) from the original SARS-CoV-2 strain (D614) were generated using previously described methods.^78,79^ To determine the binding titers of serum samples to spike or NP, we immobilized 50 ng/well of S2P trimer or nucleocapsid protein onto ELISA plates and incubated them overnight at 4°C. Subsequently, the ELISA plates were blocked with 300 μl blocking buffer (1% BSA and 10% bovine calf serum (BCS) (Sigma)) in PBS at 37°C for 2 hours. Afterwards, serum samples were serially diluted by 5-fold from 100× using dilution buffer (1% BSA and 20% BCS in PBS) and incubated in the ELISA plates at 37°C for 1 h. After incubation, 10,000-fold diluted Peroxidase AffiniPure goat anti-human IgG (H+L) antibody (Jackson ImmunoResearch) was added and incubated for 1 h at 37°C. The plates were washed between each step with PBST (0.5% Tween-20 in PBS). Finally, the TMB substrate (Sigma) was added and incubated before the reaction was stopped using 1 M sulfuric acid. Absorbance was measured at 450 nm. EC50 values was calculated as the dilutions at which the OD450 readings reached half of the maximal using GraphPad Prism v.9.2.

### PBMCs with peptide stimulation for single-cell analysis

A cryovial that contains frozen PBMCs (10^6^ cells/vial) from each of 11 COVID-19 patients was thawed in 37°C water bath. Immediately after, thawed cells were transferred to 15-mL Falcon tube that contains 10 mL of cold culture medium. After centrifugation at 150g for 5 min, the cell pellets were resuspended in complete RPMI + 10% heat-inactivated FCS medium and plated onto 12-well plate at a concentration of 5 x 10^6^ cells/3 mL/well. Then a cocktail of peptides (6 peptides; FAQDGNAAI, LMIERFVSL, ILHCANFNV, VLNDILSRL, VVFLHVTYV, and KLSYGIATV) was added at a concentration of 5 μg/mL for 16hr in 5% CO2 37oC incubator. After 16-hour incubation, the cells were washed 3 times and incubated for 10 minutes with Fc receptor block (BioLegend, 422302). Afterwards, each sample was stained with a total of 6 HLA-A2^+^/peptide fluorescent tetramer (Tet) and pentamer (Pent) for 10 min (**Figure S2A**). After cell washing at twice, each sample was stained with cell hashing antibodies for 20 min, respectively (**Table S6**) and then washed at three times and finally pooled. The combined cells were stained with a cocktail of CITE-seq and anti-human CD8 FACS antibodies for 30 minutes (**Table S7**). After cell washing at twice, viable Tet/Pent^+^ CD8^+^ T cells were sorted by FACS Aria II (**Figure S2B**).

### Single-cell analysis

The scRNA-Seq libraries were prepared using the 10x Chromium single-cell 3’ reagent kits (v3.1 Chemistry), per manufacturer’s instructions. In brief, after cell sorting, single-cell suspensions were loaded into the Chromium controller to make nanoliterscale droplets with uniquely barcoded 3’ gel beads called GEMs (gel bead-in emulsions). After GEM-RT, GEMs were cleaned up by Dyna beads MyOne Silane beads (Thermo Fisher Scientific, 37002D). According to the manufacturer’s instructions, the cDNA was amplified and size-selected by SPRI-beads (Beckman Coulter, B23317). Finally, the 3’ transcript and feature barcode libraries were pooled and sequenced with the Illumina NovaSeq 6000 sequencing system.

### Data processing of scRNA-seq

FASTQ files were processed by Cell Ranger v.6.1.2 (10x Genomics) software using human GRCh38 (GENCODE v32/Ensembl98) genome as a reference. The data analyzed in R using the Seurat v5 package were merged and integrated using an anchor-based single-cell data integration method in Seurat. Cells with mitochondrial RNA content greater than 10% were excluded. Variable feature genes were set at 2000 genes. Once the data sets were integrated, the data were input into a principal component analysis (PCA) based on variable genes. The same principal components were used to generate the Uniform Manifold Approximation and Projections (UMAPs). Clusters were identified using shared nearest neighbor–based (SNN-based) clustering. For the clustering analysis, the function RunUMAP, FindClusters, and Find-Neighbors in Seurat were used.

### Single cell trajectory

The Seurat object including 13 clusters that were characterized by unsupervised clustering in **Figure 4B**, was employed for the single-cell trajectory analysis by using Monocle3, which is available at (https://cole-trapnell-lab.github.io/monocle3/). The data from the Seurat object (RNA expression, metadata, and gene annotation) were converted into Monocle3 with a Seurat function “ProjectDim()”. Then, a “newCellDataSet()” with monocle3 was created. The PCA, UMAP, and cell clusters from the Seurat were embedded into the newCellDataset. Afterwards, using the “learn_graph()” function, single cells were partitioned in a trajectory graph. Then, based on the pseudotime (root node: the T_naive_ cluster), the cells were ordered with “order_cells()” function and visualized.

## Data and code availability

This paper does not report original code. Any additional information required to reanalyze the data reported in this paper is available from the corresponding authors upon request.

## Acknowledgements

We thank Dr. David D. Ho and Dr. Donna Farber for their valuable comments and supports. The study was supported by funding from NIH: K23 AI171263 (to L.J.P.), and K24 AI155230 (to M.T.Y.), and the institutional internal funding to M.T.

## Contributions

K.M., S.I., and M.T. conceived and designed the study. K.M., S.I., J.H., L.L., Q.W. and M.T. performed experiments. L.F.L. and S.I. validated tetramers and pentamers. K.M. performed cell sorting and single cell RNA-sequencing. K.M., Y.Q., and Z.S. performed computational analyses. J.S., C.R., M.L.M., L.J.P. and M.Y. collected human samples and/or data. K.M., S.I., L.L., C.G. J.G.A.C-d-R, Z.S., M.Y., and M.T. analyzed the data. L.J.P, M.Y., and M.T. funded the study. K.M. and M.T. wrote the manuscript; all authors edited and approved the paper.

## Corresponding authors

*Correspondence to Kazuya Masuda and Moriya Tsuji

## References

1. Josephson, A., Kilic, T., and Michler, J.D. (2021). Socioeconomic impacts of COVID-19 in low-income countries. Nat. Hum. Behav. 5, 557–565. 10.1038/s41562-021-01096-7.

2. Carabelli, A.M., Peacock, T.P., Thorne, L.G., Harvey, W.T., Hughes, J., COVID-19 Genomics UK Consortium, De Silva, T.I., Peacock, S.J., Barclay, W.S., De Silva, T.I., et al. (2023). SARS-CoV-2 variant biology: immune escape, transmission and fitness. Nat. Rev. Microbiol. 10.1038/s41579-022-00841-7.

3. Cox, M., Peacock, T.P., Harvey, W.T., Hughes, J., Wright, D.W., COVID-19 Genomics UK (COG-UK) Consortium, Willett, B.J., Thomson, E., Gupta, R.K., Peacock, S.J., et al. (2023). SARS-CoV-2 variant evasion of monoclonal antibodies based on in vitro studies. Nat. Rev. Microbiol. 21, 112–124. 10.1038/s41579-022-00809-7.

4. Taylor, P.C., Adams, A.C., Hufford, M.M., de la Torre, I., Winthrop, K., and Gottlieb, R.L. (2021). Neutralizing monoclonal antibodies for treatment of COVID-19. Nat. Rev. Immunol. 21, 382–393. 10.1038/s41577-021-00542-x.

5. Jena, D., Ghosh, A., Jha, A., Prasad, P., and Raghav, S.K. (2024). Impact of vaccination on SARS-CoV-2 evolution and immune escape variants. Vaccine 42, 126153. 10.1016/j.vaccine.2024.07.054.

6. Liu, J., Pan, X., Zhang, S., Li, M., Ma, K., Fan, C., Lv, Y., Guan, X., Yang, Y., Ye, X., et al. (2023). Efficacy and safety of Paxlovid in severe adult patients with SARS-Cov-2 infection: a multicenter randomized controlled study. Lancet Reg. Health West. Pac. 33, 100694. 10.1016/j.lanwpc.2023.100694.

7. Butler, C.C., Hobbs, F.D.R., Gbinigie, O.A., Rahman, N.M., Hayward, G., Richards, D.B., Dorward, J., Lowe, D.M., Standing, J.F., Breuer, J., et al. (2023). Molnupiravir plus usual care versus usual care alone as early treatment for adults with COVID-19 at increased risk of adverse outcomes (PANORAMIC): an open-label, platform-adaptive randomised controlled trial. Lancet Lond. Engl. 401, 281–293. 10.1016/S0140-6736(22)02597-1.

8. WHO Solidarity Trial Consortium, Pan, H., Peto, R., Henao-Restrepo, A.-M., Preziosi, M.-P., Sathiyamoorthy, V., Abdool Karim, Q., Alejandria, M.M., Hernández García, C., Kieny, M.-P., et al. (2021). Repurposed Antiviral Drugs for Covid-19 - Interim WHO Solidarity Trial Results. N. Engl. J. Med. 384, 497–511. 10.1056/NEJMoa2023184.

9. Cai, M., Xie, Y., Topol, E.J., and Al-Aly, Z. (2024). Three-year outcomes of post-acute sequelae of COVID-19. Nat. Med. 30, 1564–1573. 10.1038/s41591-024-02987-8.

10. Xu, Q., Milanez-Almeida, P., Martins, A.J., Radtke, A.J., Hoehn, K.B., Oguz, C., Chen, J., Liu, C., Tang, J., Grubbs, G., et al. (2023). Adaptive immune responses to SARS-CoV-2 persist in the pharyngeal lymphoid tissue of children. Nat. Immunol. 24, 186–199. 10.1038/s41590-022-01367-z.

11. Guo, L., Zhang, Q., Gu, X., Ren, L., Huang, T., Li, Y., Zhang, H., Liu, Y., Zhong, J., Wang, X., et al. (2024). Durability and cross-reactive immune memory to SARS-CoV-2 in individuals 2 years after recovery from COVID-19: a longitudinal cohort study. Lancet Microbe 5, e24–e33. 10.1016/S2666-5247(23)00255-0.

12. Choy, C., Chen, J., Li, J., Gallagher, D.T., Lu, J., Wu, D., Zou, A., Hemani, H., Baptiste, B.A., Wichmann, E., et al. (2023). SARS-CoV-2 infection establishes a stable and age-independent CD8+ T cell response against a dominant nucleocapsid epitope using restricted T cell receptors. Nat. Commun. 14, 6725. 10.1038/s41467-023-42430-z.

13. Meyer, S., Blaas, I., Bollineni, R.C., Delic-Sarac, M., Tran, T.T., Knetter, C., Dai, K.-Z., Madssen, T.S., Vaage, J.T., Gustavsen, A., et al. (2023). Prevalent and immunodominant CD8 T cell epitopes are conserved in SARS-CoV-2 variants. Cell Rep. 42, 111995. 10.1016/j.celrep.2023.111995.

14. Minervina, A.A., Pogorelyy, M.V., Kirk, A.M., Crawford, J.C., Allen, E.K., Chou, C.-H., Mettelman, R.C., Allison, K.J., Lin, C.-Y., Brice, D.C., et al. (2022). SARS-CoV-2 antigen exposure history shapes phenotypes and specificity of memory CD8+ T cells. Nat. Immunol. 23, 781–790. 10.1038/s41590-022-01184-4.

15. Gao, Y., Cai, C., Grifoni, A., Müller, T.R., Niessl, J., Olofsson, A., Humbert, M., Hansson, L., Österborg, A., Bergman, P., et al. (2022). Ancestral SARS-CoV-2-specific T cells cross-recognize the Omicron variant. Nat. Med. 28, 472–476. 10.1038/s41591-022-01700-x.

16. Dolton, G., Rius, C., Hasan, M.S., Wall, A., Szomolay, B., Behiry, E., Whalley, T., Southgate, J., Fuller, A., COVID-19 Genomics UK (COG-UK) consortium, et al. (2022). Emergence of immune escape at dominant SARS-CoV-2 killer T cell epitope. Cell 185, 2936-2951.e19. 10.1016/j.cell.2022.07.002.

17. Schulien, I., Kemming, J., Oberhardt, V., Wild, K., Seidel, L.M., Killmer, S., Sagar, null, Daul, F., Salvat Lago, M., Decker, A., et al. (2021). Characterization of pre-existing and induced SARS-CoV-2-specific CD8+ T cells. Nat. Med. 27, 78–85. 10.1038/s41591-020-01143-2.

18. Wagner, K.I., Mateyka, L.M., Jarosch, S., Grass, V., Weber, S., Schober, K., Hammel, M., Burrell, T., Kalali, B., Poppert, H., et al. (2022). Recruitment of highly cytotoxic CD8+ T cell receptors in mild SARS-CoV-2 infection. Cell Rep. 38, 110214. 10.1016/j.celrep.2021.110214.

19. Nesterenko, P.A., McLaughlin, J., Tsai, B.L., Burton Sojo, G., Cheng, D., Zhao, D., Mao, Z., Bangayan, N.J., Obusan, M.B., Su, Y., et al. (2021). HLA-A∗02:01 restricted T cell receptors against the highly conserved SARS-CoV-2 polymerase cross-react with human coronaviruses. Cell Rep. 37, 110167. 10.1016/j.celrep.2021.110167.

20. Zhang, H., Deng, S., Ren, L., Zheng, P., Hu, X., Jin, T., and Tan, X. (2021). Profiling CD8+ T cell epitopes of COVID-19 convalescents reveals reduced cellular immune responses to SARS-CoV-2 variants. Cell Rep. 36, 109708. 10.1016/j.celrep.2021.109708.

21. Kusnadi, A., Ramírez-Suástegui, C., Fajardo, V., Chee, S.J., Meckiff, B.J., Simon, H., Pelosi, E., Seumois, G., Ay, F., Vijayanand, P., et al. (2021). Severely ill COVID-19 patients display impaired exhaustion features in SARS-CoV-2-reactive CD8+ T cells. Sci. Immunol. 6, eabe4782. 10.1126/sciimmunol.abe4782.

22. Mallajosyula, V., Ganjavi, C., Chakraborty, S., McSween, A.M., Pavlovitch-Bedzyk, A.J., Wilhelmy, J., Nau, A., Manohar, M., Nadeau, K.C., and Davis, M.M. (2021). CD8+ T cells specific for conserved coronavirus epitopes correlate with milder disease in COVID-19 patients. Sci. Immunol. 6, eabg5669. 10.1126/sciimmunol.abg5669.

23. Lineburg, K.E., Grant, E.J., Swaminathan, S., Chatzileontiadou, D.S.M., Szeto, C., Sloane, H., Panikkar, A., Raju, J., Crooks, P., Rehan, S., et al. (2021). CD8+ T cells specific for an immunodominant SARS-CoV-2 nucleocapsid epitope cross-react with selective seasonal coronaviruses. Immunity 54, 1055–1065.e5. 10.1016/j.immuni.2021.04.006.

24. Neidleman, J., Luo, X., George, A.F., McGregor, M., Yang, J., Yun, C., Murray, V., Gill, G., Greene, W.C., Vasquez, J., et al. (2021). Distinctive features of SARS-CoV-2-specific T cells predict recovery from severe COVID-19. Cell Rep. 36, 109414. 10.1016/j.celrep.2021.109414.

25. Sette, A., and Crotty, S. (2021). Adaptive immunity to SARS-CoV-2 and COVID-19. Cell 184, 861–880. 10.1016/j.cell.2021.01.007.

26. Ferretti, A.P., Kula, T., Wang, Y., Nguyen, D.M.V., Weinheimer, A., Dunlap, G.S., Xu, Q., Nabilsi, N., Perullo, C.R., Cristofaro, A.W., et al. (2020). Unbiased Screens Show CD8+ T Cells of COVID-19 Patients Recognize Shared Epitopes in SARS-CoV-2 that Largely Reside outside the Spike Protein. Immunity 53, 1095–1107.e3. 10.1016/j.immuni.2020.10.006.

27. Shomuradova, A.S., Vagida, M.S., Sheetikov, S.A., Zornikova, K.V., Kiryukhin, D., Titov, A., Peshkova, I.O., Khmelevskaya, A., Dianov, D.V., Malasheva, M., et al. (2020). SARS-CoV-2 Epitopes Are Recognized by a Public and Diverse Repertoire of Human T Cell Receptors. Immunity 53, 1245–1257.e5. 10.1016/j.immuni.2020.11.004.

28. Sekine, T., Perez-Potti, A., Rivera-Ballesteros, O., Strålin, K., Gorin, J.-B., Olsson, A., Llewellyn-Lacey, S., Kamal, H., Bogdanovic, G., Muschiol, S., et al. (2020). Robust T Cell Immunity in Convalescent Individuals with Asymptomatic or Mild COVID-19. Cell 183, 158–168.e14. 10.1016/j.cell.2020.08.017.

29. Grifoni, A., Weiskopf, D., Ramirez, S.I., Mateus, J., Dan, J.M., Moderbacher, C.R., Rawlings, S.A., Sutherland, A., Premkumar, L., Jadi, R.S., et al. (2020). Targets of T Cell Responses to SARS-CoV-2 Coronavirus in Humans with COVID-19 Disease and Unexposed Individuals. Cell 181, 1489–1501.e15. 10.1016/j.cell.2020.05.015.

30. Lopes-Ribeiro, Á., Oliveira, P. de M., Retes, H.M., Barbosa-Stancioli, E.F., da Fonseca, F.G., Tsuji, M., and Coelho-Dos-Reis, J.G.A. (2023). Surveillance of SARS-CoV-2 immunogenicity: loss of immunodominant HLA-A*02-restricted epitopes that activate CD8+ T cells. Front. Immunol. 14, 1229712. 10.3389/fimmu.2023.1229712.

31. Zonozi, R., Walters, L.C., Shulkin, A., Naranbhai, V., Nithagon, P., Sauvage, G., Kaeske, C., Cosgrove, K., Nathan, A., Tano-Menka, R., et al. T cell responses to SARS-CoV-2 infection and vaccination are elevated in B cell deficiency and reduce risk of severe COVID-19. Sci. Transl. Med.

32. Fumagalli, V., Ravà, M., Marotta, D., Di Lucia, P., Bono, E.B., Giustini, L., De Leo, F., Casalgrandi, M., Monteleone, E., Mouro, V., et al. (2024). Antibody-independent protection against heterologous SARS-CoV-2 challenge conferred by prior infection or vaccination. Nat. Immunol. 25, 633–643. 10.1038/s41590-024-01787-z.

33. Wu, Y., and Chen, Y. Reduction and Functional Exhaustion of T Cells in Patients With Coronavirus Disease 2019 (COVID-19).

34. Arieta, C.M., Xie, Y.J., Rothenberg, D.A., Diao, H., Harjanto, D., Meda, S., Marquart, K., Koenitzer, B., Sciuto, T.E., Lobo, A., et al. (2023). The T-cell-directed vaccine BNT162b4 encoding conserved non-spike antigens protects animals from severe SARS-CoV-2 infection. Cell 186, 2392–2409.e21. 10.1016/j.cell.2023.04.007.

35. Adamo, S., Michler, J., Zurbuchen, Y., Cervia, C., Taeschler, P., Raeber, M.E., Baghai Sain, S., Nilsson, J., Moor, A.E., and Boyman, O. (2022). Signature of long-lived memory CD8+ T cells in acute SARS-CoV-2 infection. Nature 602, 148–155. 10.1038/s41586-021-04280-x.

36. Zhang, B., Upadhyay, R., Hao, Y., Samanovic, M.I., Herati, R.S., Blair, J.D., Axelrad, J., Mulligan, M.J., Littman, D.R., and Satija, R. (2023). Multimodal single-cell datasets characterize antigen-specific CD8+ T cells across SARS-CoV-2 vaccination and infection. Nat. Immunol. 24, 1725–1734. 10.1038/s41590-023-01608-9.

37. Unterman, A., Sumida, T.S., Nouri, N., Yan, X., Zhao, A.Y., Gasque, V., Schupp, J.C., Asashima, H., Liu, Y., Cosme, C., et al. (2022). Single-cell multi-omics reveals dyssynchrony of the innate and adaptive immune system in progressive COVID-19. Nat. Commun. 13, 440. 10.1038/s41467-021-27716-4.

38. Zhang, J.-Y., Wang, X.-M., Xing, X., Xu, Z., Zhang, C., Song, J.-W., Fan, X., Xia, P., Fu, J.-L., Wang, S.-Y., et al. (2020). Single-cell landscape of immunological responses in patients with COVID-19. Nat. Immunol. 21, 1107–1118. 10.1038/s41590-020-0762-x.

39. Ellis, J.M., Henson, V., Slack, R., Ng, J., Hartzman, R.J., and Katovich Hurley, C. (2000). Frequencies of HLA-A2 alleles in five U.S. population groups. Predominance Of A*02011 and identification of HLA-A*0231. Hum. Immunol. 61, 334–340. 10.1016/s0198-8859(99)00155-x.

40. Augusto, D.G., Murdolo, L.D., Chatzileontiadou, D.S.M., Sabatino, J.J., Yusufali, T., Peyser, N.D., Butcher, X., Kizer, K., Guthrie, K., Murray, V.W., et al. (2023). A common allele of HLA is associated with asymptomatic SARS-CoV-2 infection. Nature 620, 128–136. 10.1038/s41586-023-06331-x.

41. WHO Working Group on the Clinical Characterisation and Management of COVID-19 infection (2020). A minimal common outcome measure set for COVID-19 clinical research. Lancet Infect. Dis. 20, e192–e197. 10.1016/S1473-3099(20)30483-7.

42. Chia, W.N., Zhu, F., Ong, S.W.X., Young, B.E., Fong, S.-W., Le Bert, N., Tan, C.W., Tiu, C., Zhang, J., Tan, S.Y., et al. (2021). Dynamics of SARS-CoV-2 neutralising antibody responses and duration of immunity: a longitudinal study. Lancet Microbe 2, e240–e249. 10.1016/S2666-5247(21)00025-2.

43. Seow, J., Graham, C., Merrick, B., Acors, S., Pickering, S., Steel, K.J.A., Hemmings, O., O’Byrne, A., Kouphou, N., Galao, R.P., et al. (2020). Longitudinal observation and decline of neutralizing antibody responses in the three months following SARS-CoV-2 infection in humans. Nat. Microbiol. 5, 1598–1607. 10.1038/s41564-020-00813-8.

44. Zuo, J., Dowell, A.C., Pearce, H., Verma, K., Long, H.M., Begum, J., Aiano, F., Amin-Chowdhury, Z., Hoschler, K., Brooks, T., et al. (2021). Robust SARS-CoV-2-specific T cell immunity is maintained at 6 months following primary infection. Nat. Immunol. 22, 620–626. 10.1038/s41590-021-00902-8.

45. Khoury, D.S., Cromer, D., Reynaldi, A., Schlub, T.E., Wheatley, A.K., Juno, J.A., Subbarao, K., Kent, S.J., Triccas, J.A., and Davenport, M.P. (2021). Neutralizing antibody levels are highly predictive of immune protection from symptomatic SARS-CoV-2 infection. Nat. Med. 27, 1205–1211. 10.1038/s41591-021-01377-8.

46. Rydyznski Moderbacher, C., Ramirez, S.I., Dan, J.M., Grifoni, A., Hastie, K.M., Weiskopf, D., Belanger, S., Abbott, R.K., Kim, C., Choi, J., et al. (2020). Antigen-Specific Adaptive Immunity to SARS-CoV-2 in Acute COVID-19 and Associations with Age and Disease Severity. Cell 183, 996–1012.e19. 10.1016/j.cell.2020.09.038.

47. Cromer, D., Steain, M., Reynaldi, A., Schlub, T.E., Wheatley, A.K., Juno, J.A., Kent, S.J., Triccas, J.A., Khoury, D.S., and Davenport, M.P. (2022). Neutralising antibody titres as predictors of protection against SARS-CoV-2 variants and the impact of boosting: a meta-analysis. Lancet Microbe 3, e52–e61. 10.1016/S2666-5247(21)00267-6.

48. Garcia-Beltran, W.F., Lam, E.C., Astudillo, M.G., Yang, D., Miller, T.E., Feldman, J., Hauser, B.M., Caradonna, T.M., Clayton, K.L., Nitido, A.D., et al. (2021). COVID-19-neutralizing antibodies predict disease severity and survival. Cell 184, 476–488.e11. 10.1016/j.cell.2020.12.015.

49. Lockhart, A., Mucida, D., and Bilate, A.M. (2024). Intraepithelial Lymphocytes of the Intestine. Annu. Rev. Immunol. 42, 289–316. 10.1146/annurev-immunol-090222-100246.

50. Fergusson, J.R., Smith, K.E., Fleming, V.M., Rajoriya, N., Newell, E.W., Simmons, R., Marchi, E., Björkander, S., Kang, Y.-H., Swadling, L., et al. (2014). CD161 defines a transcriptional and functional phenotype across distinct human T cell lineages. Cell Rep. 9, 1075–1088. 10.1016/j.celrep.2014.09.045.

51. Castellino, F., Huang, A.Y., Altan-Bonnet, G., Stoll, S., Scheinecker, C., and Germain, R.N. (2006). Chemokines enhance immunity by guiding naive CD8+ T cells to sites of CD4+ T cell–dendritic cell interaction. Nature 440, 890–895. 10.1038/nature04651.

52. Yang, C.Y., Best, J.A., Knell, J., Yang, E., Sheridan, A.D., Jesionek, A.K., Li, H.S., Rivera, R.R., Lind, K.C., D’Cruz, L.M., et al. (2011). The transcriptional regulators Id2 and Id3 control the formation of distinct memory CD8+ T cell subsets. Nat. Immunol. 12, 1221– 1229. 10.1038/ni.2158.

53. Guo, L., Liu, X., and Su, X. (2023). The role of TEMRA cell-mediated immune senescence in the development and treatment of HIV disease. Front. Immunol. 14, 1284293. 10.3389/fimmu.2023.1284293.

54. Du, J., Wei, L., Li, G., Hua, M., Sun, Y., Wang, D., Han, K., Yan, Y., Song, C., Song, R., et al. (2021). Persistent High Percentage of HLA-DR+CD38high CD8+ T Cells Associated With Immune Disorder and Disease Severity of COVID-19. Front. Immunol. 12, 735125. 10.3389/fimmu.2021.735125.

55. Leonard, B., Starrett, G.J., Maurer, M.J., Oberg, A.L., Van Bockstal, M., Van Dorpe, J., De Wever, O., Helleman, J., Sieuwerts, A.M., Berns, E.M.J.J., et al. (2016). APOBEC3G Expression Correlates with T-Cell Infiltration and Improved Clinical Outcomes in High-grade Serous Ovarian Carcinoma. Clin. Cancer Res. 22, 4746–4755. 10.1158/1078-0432.CCR-15-2910.

56. Fournier, P.G.J., Juárez, P., Jiang, G., Clines, G.A., Niewolna, M., Kim, H.S., Walton, H.W., Peng, X.H., Liu, Y., Mohammad, K.S., et al. (2015). The TGF-β Signaling Regulator PMEPA1 Suppresses Prostate Cancer Metastases to Bone. Cancer Cell 27, 809–821. 10.1016/j.ccell.2015.04.009.

57. Mei, J., Liu, Y., Yu, X., Hao, L., Ma, T., Zhan, Q., Zhang, Y., and Zhu, Y. (2021). YWHAZ interacts with DAAM1 to promote cell migration in breast cancer. Cell Death Discov. 7, 221. 10.1038/s41420-021-00609-7.

58. Moss, P. (2022). The T cell immune response against SARS-CoV-2. Nat. Immunol. 23, 186–193. 10.1038/s41590-021-01122-w.

59. Markov, N.S., Ren, Z., Senkow, K.J., Grant, R.A., Gao, C.A., Malsin, E.S., Sichizya, L., Kihshen, H., Helmin, K.A., Jovisic, M., et al. (2024). Distinctive evolution of alveolar T cell responses is associated with clinical outcomes in unvaccinated patients with SARS-CoV-2 pneumonia. Nat. Immunol. 25, 1607–1622. 10.1038/s41590-024-01914-w.

60. Konduri, V., Oyewole-Said, D., Vazquez-Perez, J., Weldon, S.A., Halpert, M.M., Levitt, J.M., and Decker, W.K. (2021). CD8+CD161+ T-Cells: Cytotoxic Memory Cells With High Therapeutic Potential. Front. Immunol. 11, 613204. 10.3389/fimmu.2020.613204.

61. Liu, Y., Wang, W., Zhu, P., Cheng, X., Wu, M., Zhang, H., Chen, Y., Chen, Y., Liang, Z., Wu, X., et al. (2023). Increased Non-MAIT CD161+CD8+ T Cells Display Pathogenic Potential in Chronic HBV Infection. Cell. Mol. Gastroenterol. Hepatol. 15, 1181–1198. 10.1016/j.jcmgh.2023.02.001.

62. Flament, H., Rouland, M., Beaudoin, L., Toubal, A., Bertrand, L., Lebourgeois, S., Rousseau, C., Soulard, P., Gouda, Z., Cagninacci, L., et al. (2021). Outcome of SARS-CoV-2 infection is linked to MAIT cell activation and cytotoxicity. Nat. Immunol. 22, 322–335. 10.1038/s41590-021-00870-z.

63. Walker, L.J., Marrinan, E., Muenchhoff, M., Ferguson, J., Kloverpris, H., Cheroutre, H., Barnes, E., Goulder, P., and Klenerman, P. (2013). CD8αα Expression Marks Terminally Differentiated Human CD8+ T Cells Expanded in Chronic Viral Infection. Front. Immunol. 4. 10.3389/fimmu.2013.00223.

64. Vandereyken, M., James, O.J., and Swamy, M. (2020). Mechanisms of activation of innate-like intraepithelial T lymphocytes. Mucosal Immunol. 13, 721–731. 10.1038/s41385-020-0294-6.

65. Ramirez, S.I., Faraji, F., Hills, L.B., Lopez, P.G., Goodwin, B., Stacey, H.D., Sutton, H.J., Hastie, K.M., Saphire, E.O., Kim, H.J., et al. (2024). Immunological memory diversity in the human upper airway. Nature 632, 630–636. 10.1038/s41586-024-07748-8.

66. Suzuki, I., and Fink, P.J. (2000). The dual functions of Fas ligand in the regulation of peripheral CD8^+^ and CD4^+^ T cells. Proc. Natl. Acad. Sci. 97, 1707–1712. 10.1073/pnas.97.4.1707.

67. Phetsouphanh, C., Darley, D.R., Wilson, D.B., Howe, A., Munier, C.M.L., Patel, S.K., Juno, J.A., Burrell, L.M., Kent, S.J., Dore, G.J., et al. (2022). Immunological dysfunction persists for 8 months following initial mild-to-moderate SARS-CoV-2 infection. Nat. Immunol. 23, 210–216. 10.1038/s41590-021-01113-x.

68. Yin, K., Peluso, M.J., Luo, X., Thomas, R., Shin, M.-G., Neidleman, J., Andrew, A., Young, K.C., Ma, T., Hoh, R., et al. (2024). Long COVID manifests with T cell dysregulation, inflammation and an uncoordinated adaptive immune response to SARS-CoV-2. Nat. Immunol. 25, 218–225. 10.1038/s41590-023-01724-6.

69. Wang, J., Zhao, X., and Wan, Y.Y. (2023). Intricacies of TGF-β signaling in Treg and Th17 cell biology. Cell. Mol. Immunol. 20, 1002–1022. 10.1038/s41423-023-01036-7.

70. Filippi, C.M., Juedes, A.E., Oldham, J.E., Ling, E., Togher, L., Peng, Y., Flavell, R.A., and Von Herrath, M.G. (2008). Transforming Growth Factor-β Suppresses the Activation of CD8+ T-Cells When Naïve but Promotes Their Survival and Function Once Antigen Experienced. Diabetes 57, 2684–2692. 10.2337/db08-0609.

71. Ang, S.-F., Zhao, Z., Lim, L., and Manser, E. (2010). DAAM1 Is a Formin Required for Centrosome Re-Orientation during Cell Migration. PLoS ONE 5, e13064. 10.1371/journal.pone.0013064.

72. Morrot, A., Hafalla, J.C.R., Cockburn, I.A., Carvalho, L.H., and Zavala, F. (2005). IL-4 receptor expression on CD8+ T cells is required for the development of protective memory responses against liver stages of malaria parasites. J. Exp. Med. 202, 551–560. 10.1084/jem.20042463.

73. Singer, M., Wang, C., Cong, L., Marjanovic, N.D., Kowalczyk, M.S., Zhang, H., Nyman, J., Sakuishi, K., Kurtulus, S., Gennert, D., et al. (2016). A Distinct Gene Module for Dysfunction Uncoupled from Activation in Tumor-Infiltrating T Cells. Cell 166, 1500–1511.e9. 10.1016/j.cell.2016.08.052.

74. Soto-Heredero, G., Gómez De Las Heras, M.M., Escrig-Larena, J.I., and Mittelbrunn, M. (2023). Extremely Differentiated T Cell Subsets Contribute to Tissue Deterioration During Aging. Annu. Rev. Immunol. 41, 181–205. 10.1146/annurev-immunol-101721-064501.

75. Cai, Z., Kishimoto, H., Brunmark, A., Jackson, M.R., Peterson, P.A., and Sprent, J. (1997). Requirements for peptide-induced T cell receptor downregulation on naive CD8+ T cells. J. Exp. Med. 185, 641–651. 10.1084/jem.185.4.641.

76. Garner, S.F., Petrochilos, J., Brown, C.J., Cavanna, S., Chanarin, I., and Navarrete, C. (1997). A clinically significant anti-HLA-A2 detectable by extended incubation cytotoxicity and flow cytometric techniques but not by a standard NIH lymphocytotoxicity test. Immunohematology 13, 49–53.

77. Jurtz, V., Paul, S., Andreatta, M., Marcatili, P., Peters, B., and Nielsen, M. (2017). NetMHCpan-4.0: Improved Peptide-MHC Class I Interaction Predictions Integrating Eluted Ligand and Peptide Binding Affinity Data. J. Immunol. Baltim. Md 1950 199, 3360–3368. 10.4049/jimmunol.1700893.

78. Wang, Q., Iketani, S., Li, Z., Liu, L., Guo, Y., Huang, Y., Bowen, A.D., Liu, M., Wang, M., Yu, J., et al. (2023). Alarming antibody evasion properties of rising SARS-CoV-2 BQ and XBB subvariants. Cell 186, 279–286.e8. 10.1016/j.cell.2022.12.018.

79. Wang, P., Liu, L., Nair, M.S., Yin, M.T., Luo, Y., Wang, Q., Yuan, T., Mori, K., Solis, A.G., Yamashita, M., et al. (2020). SARS-CoV-2 neutralizing antibody responses are more robust in patients with severe disease. Emerg. Microbes Infect. 9, 2091–2093. 10.1080/22221751.2020.1823890.

