## Supplementary information for "Distinct CD8^+^ T-cell types Associated with COVID-19 Severity in Unvaccinated HLA-A2^+^ Patients"

**Table S1. Identification of 42 HLA-A2$^+$ COVID-19 patients out of 77 patients by genotyping or phenotyping**

**Table S2. Grading of the severity of COVID-19 patients at CUIMC**

**Table S3. Predicted HLA-A2-binding peptides from WT SARS-CoV-2 AA sequence**

**Table S4. HLA-A2-restricted CD8$^+$ T-cell responses of 24 COVID patients to specific epitopes of SARS-CoV-2, as determined by ELISpot assay**

**Table S5. Sequence homology of SARS-CoV-2 epitopes among SARS-CoV-2 variants and other strains**

**Table S6. Cell hashtag list**

**Table S7. CITE-seq antibody list**

**Table S8. List of top 30 genes in $T_{IEL\text{-}like}$ and C1 cluster by differentially expressed gene (DEG) analysis, respectively**

### Figure S1

A

| Subject ID# | Genotyping by ARC | NP | S trimer | NP Ab Response | S trimer Response | Severity |
|---|---|---|---|---|---|---|
| C0002 | | 482529 | 2522 | Strong | Modest | Moderate |
| C0006 | | 1534 | 9050 | Modest | Modest | Moderate |
| C0008 | | 5475 | 1152 | Modest | Modest | Mild |
| C0009 | | 4605 | 2184 | Modest | Modest | Mild |
| C0011 | | 2445 | 2060 | Modest | Modest | Mild |
| C0017 | | 100 | 230 | Weak | Weak | Mild |
| C0018 | | 1883 | 4261 | Modest | Modest | Mild |
| C0022 | | 100 | 331 | Weak | Weak | Mild |
| C0028 | | 1526 | 810 | Modest | Weak | Mild |
| C0029 | | 2350 | 329 | Modest | Weak | Mild |
| C0030 | | 392 | 126 | Weak | Weak | Mild |
| C0032 | | 955 | 117 | Weak | Weak | Mild |
| C0034 | | 100 | 100 | Weak | Weak | Mild |
| C0035 | | 344 | 133 | Weak | Weak | Mild |
| C0039 | | 100 | 100 | Weak | Weak | Mild |
| C0041 | | 1119 | 363 | Modest | Weak | Mild |
| C0042 | | 692 | 246 | Weak | Weak | Mild |
| C0043 | | 393 | 977 | Weak | Weak | Mild |
| C0044 | | 1154 | 13565 | Modest | Strong | Severe |
| C0046 | | 3527 | 1409 | Modest | Modest | Moderate |
| C0047 | A2+ | 344 | 858 | Weak | Weak | Mild |
| C0051 | | 655 | 387 | Weak | Weak | Mild |
| C0052 | | 119 | 248 | Weak | Weak | Mild |
| C0054 | | 100 | 100 | Weak | Weak | Mild |
| C0056 | | 3799 | 632 | Modest | Weak | Mild |
| C0069 | | 1015 | 384 | Modest | Weak | Mild |
| C0078 | | 964 | 1242 | Weak | Modest | Mild |
| C0084 | | 12698 | 17355 | Strong | Strong | Severe |
| C0086 | | 169 | 100 | Weak | Weak | Mild |
| C0088 | | 1958 | 10703 | Modest | Strong | Moderate |
| C0094 | | 100 | 221 | Weak | Weak | MIld |
| C0098 | | 481777 | 40517 | Strong | Strong | Moderate |
| C0101 | | 366 | 2024 | Weak | Modest | Severe |
| C0109 | | 256 | 382 | Weak | Weak | Moderate |
| C0112 | | 2141 | 15174 | Modest | Strong | Severe |
| C0115 | | 22840 | 8426 | Strong | Modest | Moderate |
| C0118 | | 3623 | 657 | Modest | Weak | Mild |
| C0126 | | 9983 | 2032 | Modest | Modest | Severe |
| C0129 | | 1929 | 2032 | Modest | Modest | Severe |
| C0138 | | 369 | 10162 | Weak | Strong | Severe |
| C0141 | | 7917 | 17774 | Modest | Strong | Moderate |
| C0142 | | 8551 | 1154 | Modest | Modest | Severe |

<100,"Very Weak", 100≤1000,"Weak", 1000≤10000,"Modest", >10000,"Strong

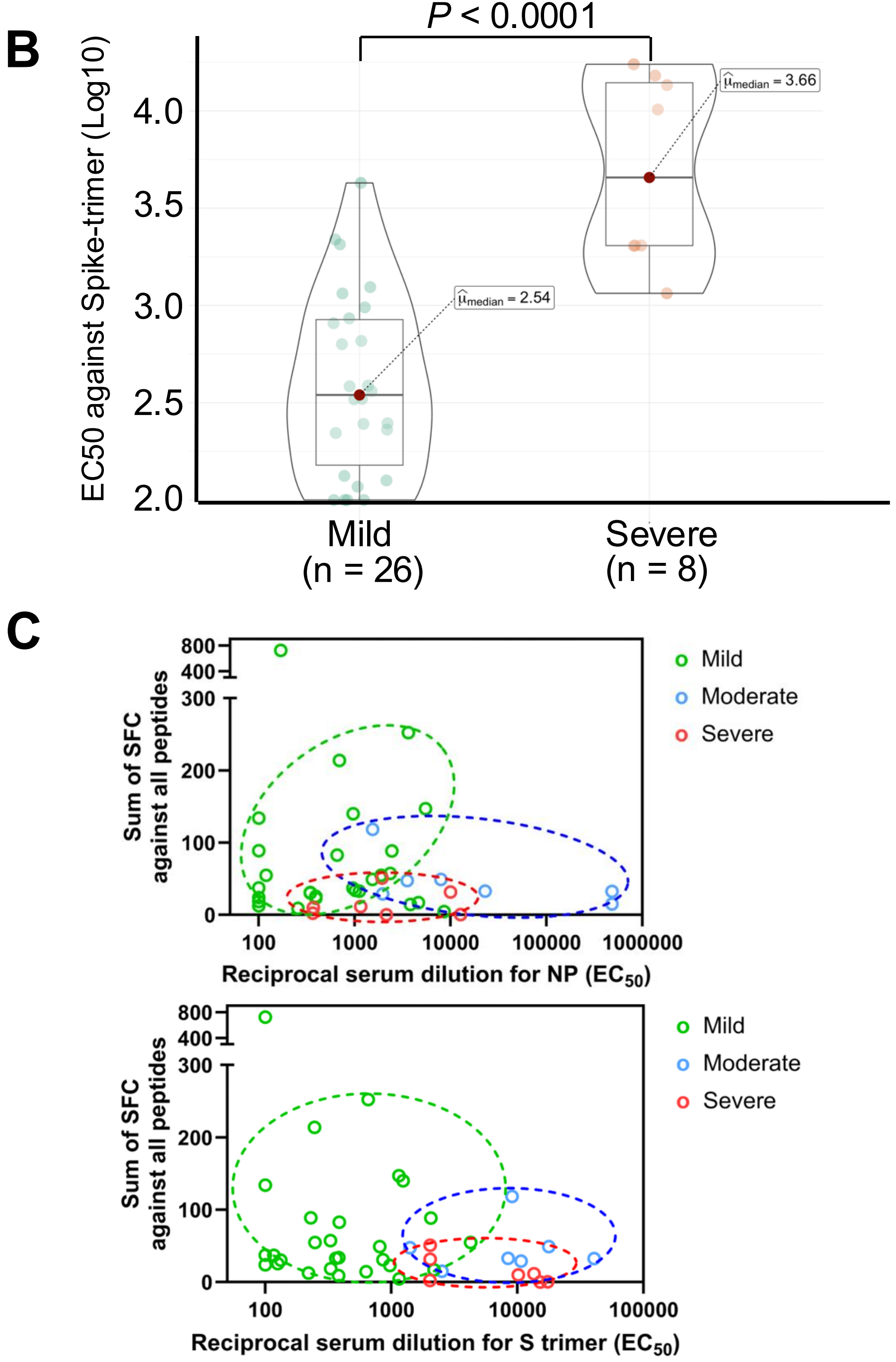

**Figure S1. Humoral and $CD8^+$ T-cell responses against SARS-CoV-2 epitopes.** (A) EC50 values of sera from 42 HLA-$A2^+$ COVID patients against N protein or Spike trimer, measured by ELISA. (B) Box and violin plots of EC50 values of sera in mild and severe COVID-19 patients and comparison of its median. A statistical value ($p < 0.0001$): Mann–Whitney U test. (C) Graph of dot plots showing EC50 values of sera (horizontal) against N protein (left) and Spike trimer (right), and a total sum of SFC against each selected peptide (vertical) from 42 HLA-$A2^+$ COVID-19 patients.

**Figure S2**

**A**

| Name of Protein | # of peptide | AA sequence | Color | Tetramer or Pentamer |
|---|---|---|---|---|
| RdR Poly | 7 | FAQDGNAAI | BV421 | Tetramer |
| RdRp | 22 | LMIERFVSL | PE | Pentamer |
| RdRp | 23 | ILHCANFNV | BV421 | Tetramer |
| S protein | 28 | VLNDILSRL | APC | Tetramer |
| S protein | 29 | VVFLHVTYV | APC | Pentamer |
| Helicase | 32 | KLSYGIATV | PE | Tetramer |

**B**

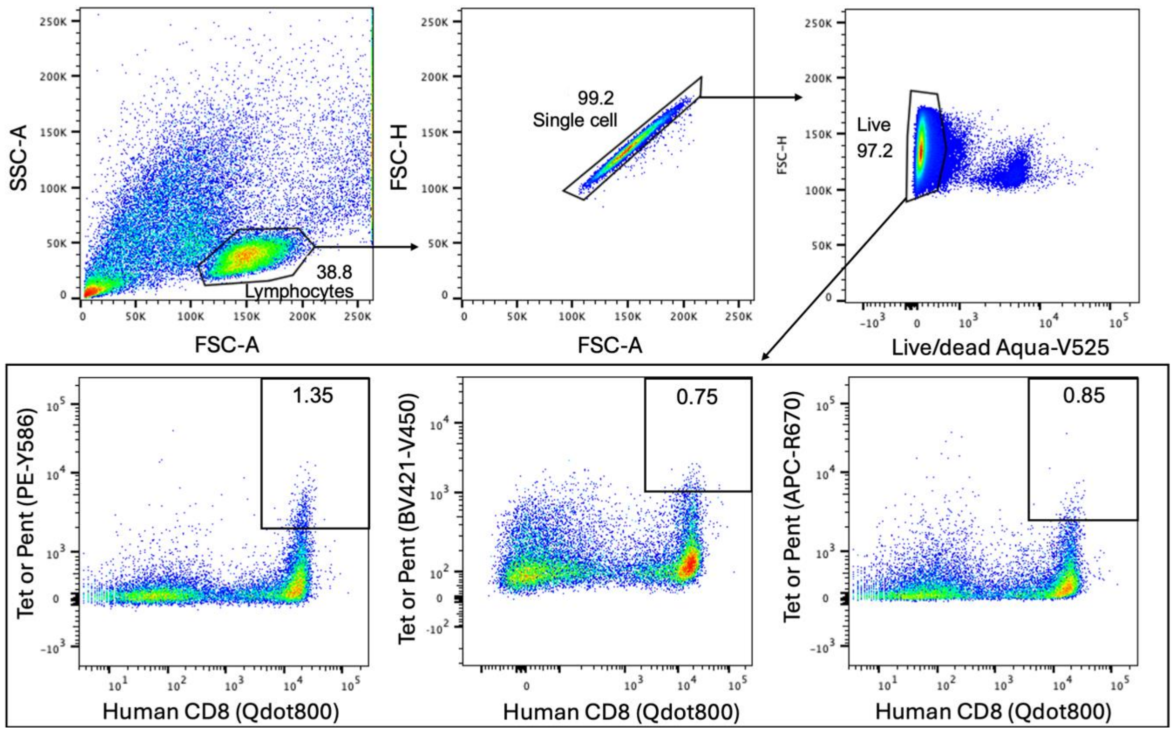

**C**

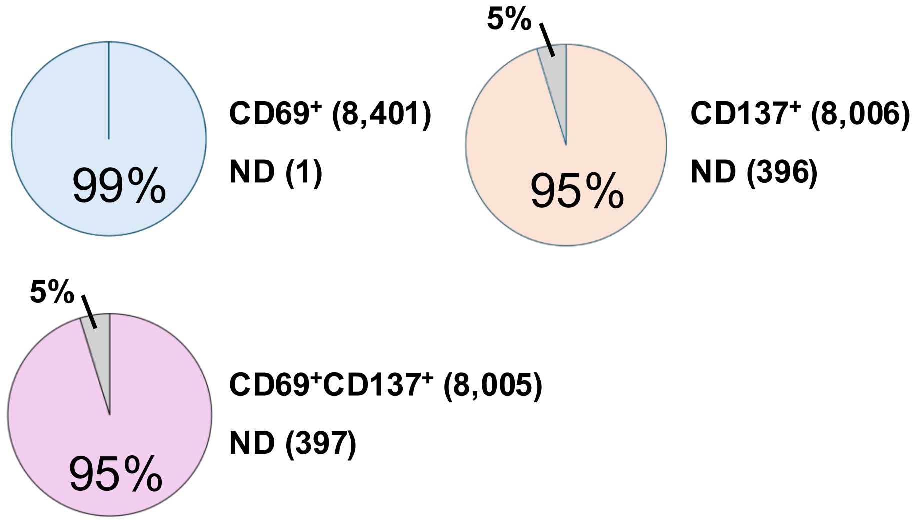

**D**

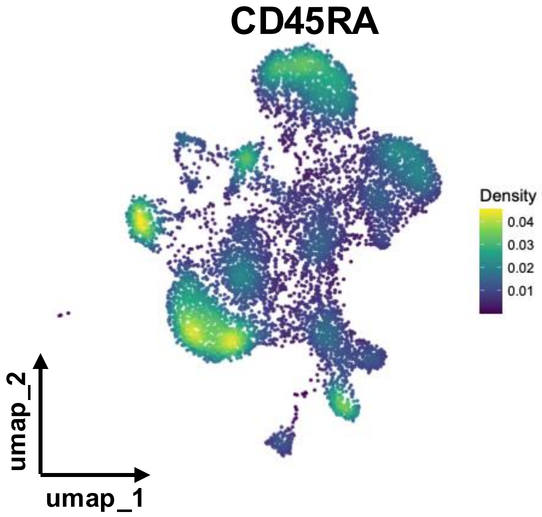

**Figure S2. Gating strategy of SARS-CoV-2 peptide loading HLA-A2$^+$ pentamer/tetramer positive CD8$^+$ T cells and CITE-seq outputs.** (A) List of fluorescent pMHC tetramers and pentamers selected from 11 immunodominant peptides. (B) FACS sorting of pMHC tetramer- and/or pentamer-positive live CD8$^+$ T cells. (C) Pie chart of CD69$^+$, CD137$^+$, or CD69$^+$CD137$^+$ cells in the UMAP. (D) Density plots of CD45RA-antigen-derived tag (ADT) expression in the UMAP.

Figure S3

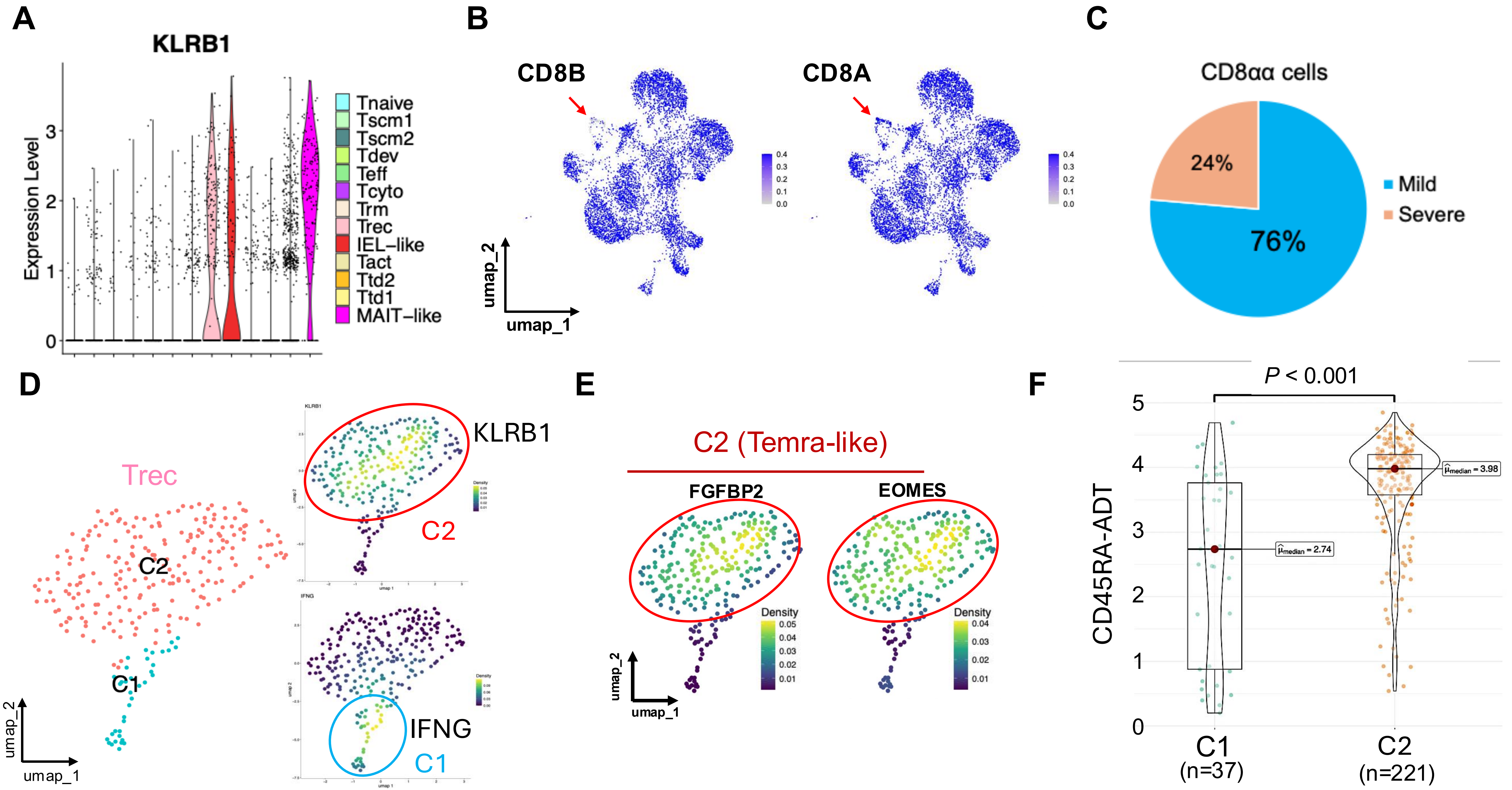
A
KLRB1
Expression Level
Tnaive
Tscm1
Tscm2
Tdev
Teff
Tcyto
Trm
Trec
IEL−like
Tact
Ttd2
Ttd1
MAIT−like
B
CD8B
CD8A
umap_2
umap_1
C
CD8αα cells
24%
76%
Mild
Severe
D
Trec
C2
C1
KLRB1
C2
IFNG
C1
E
C2 (Temra-like)
FGFBP2
EOMES
Density
F
P < 0.001
CD45RA-ADT
C1
(n=37)
C2
(n=221)

**Figure S3. Distinct features of the TIEL-like, Trec, and TMAIT-like clusters.** (A) Volcano plots of *KLRB1* gene expression (B) CD8A or CD8B gene expression in the UMAP. (C) Pie chart of CD8αα-positive cells in mild and severe COVID-19 patients. (D) Sub-clustering of UMAP plots in the $T_{rec}$ cluster (C1 and C2 clusters were colored at blue and red, respectively) (left). Density plots of *KLRB1* and *IFNG* expression in the $T_{rec}$ cluster (C1 and C2 clusters were circled at blue and red) (right). (E) Density plots of *FGFBP2* and *EOMES* expression in the $T_{rec}$ cluster (C2 clusters were circled at red). (F) Box and violin plots of CD45RA-ADT expression in the C1 and C2 clusters within the $T_{rec}$ population and comparison of its median. A statistical value ($p < 0.001$): Mann–Whitney U test.

Figure S4

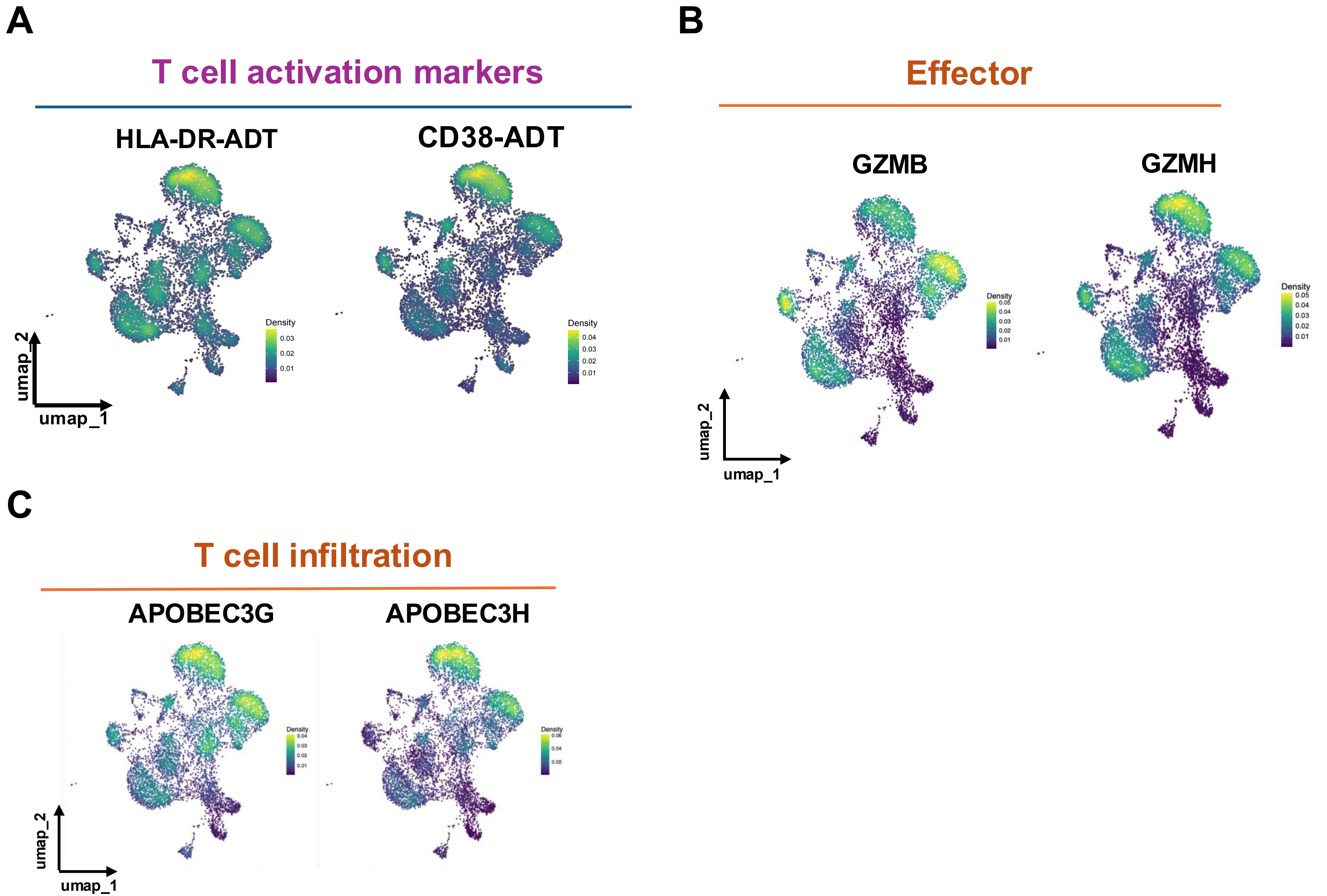
A
T cell activation markers
HLA-DR-ADT
CD38-ADT
Density
0.03
0.02
0.01
Density
0.04
0.03
0.02
0.01
umap_2
umap_1
B
Effector
GZMB
GZMH
Density
0.05
0.04
0.03
0.02
0.01
umap_2
umap_1
C
T cell infiltration
APOBEC3G
APOBEC3H
umap_2
umap_1

**Figure S4. Gene features and cell surface markers in distinct terminally differentiated CD8$^+$ T subsets.** (A) Expression of T cell activation antigen-derived tag (ADT) markers HLA-DR and CD38 in the UMAP. (B and C) Expression of effector and T cell infiltration markers in the UMAP.

**Figure S5**

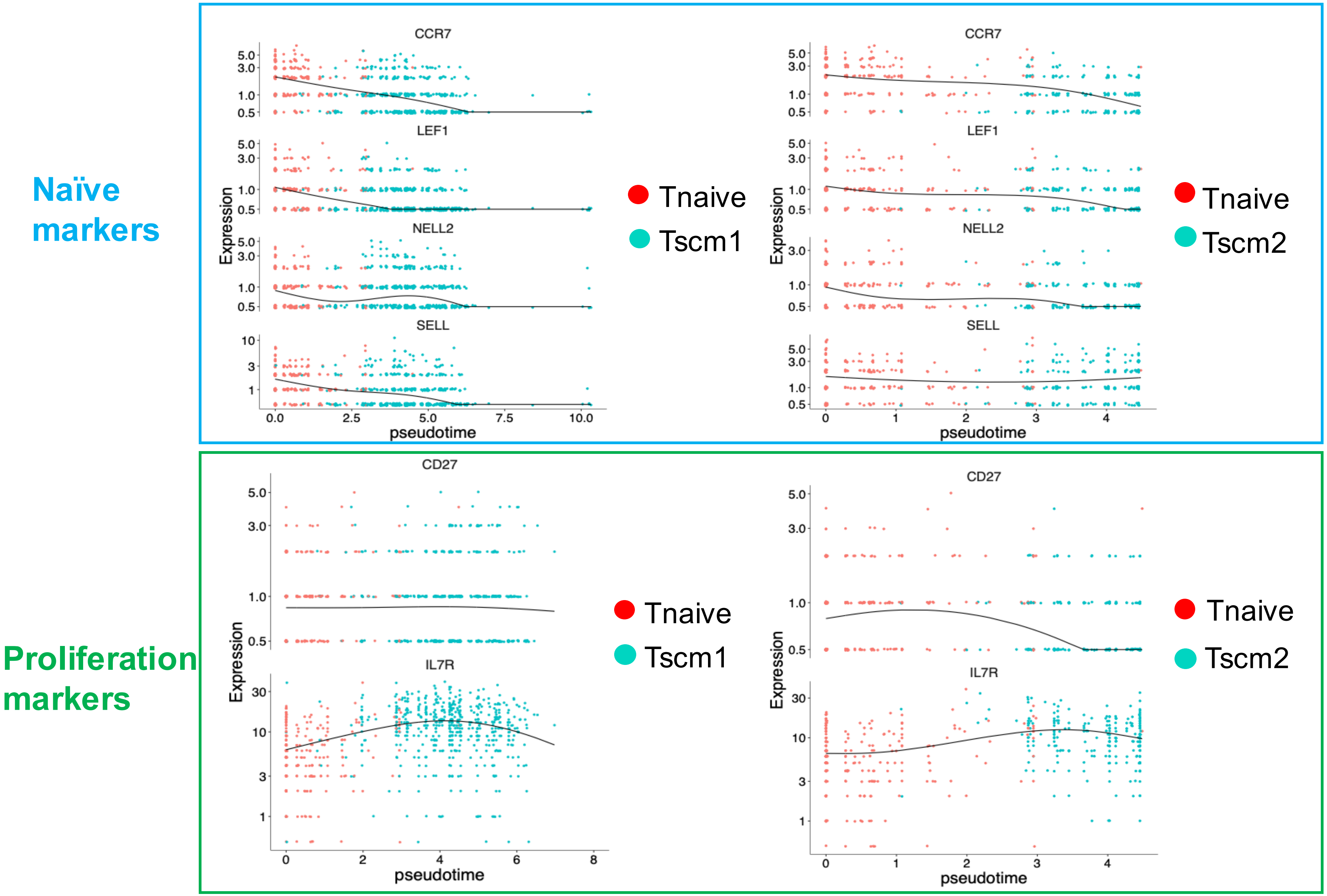

Naïve markers
CCR7
LEF1
NELL2
SELL
Expression
pseudotime
Tnaive
Tscm1
Tscm2
Proliferation markers
CD27
IL7R

**Figure S5. Relative gene expression in pseudotime of single cell trajectory.** Relative expression of *CCR7*, *LEF1*, *NELL2* and *SELL* across pseudotime in the $T_{naive}$ and $T_{scm1}$ clusters or the $T_{naive}$ and $T_{scm2}$ clusters, respectively (upper). Relative expression of *IL7R* and *CD27* across pseudotime in the $T_{naive}$ and $T_{scm1}$ clusters or the $T_{naive}$ and $T_{scm2}$ clusters, respectively (lower).

Figure S6

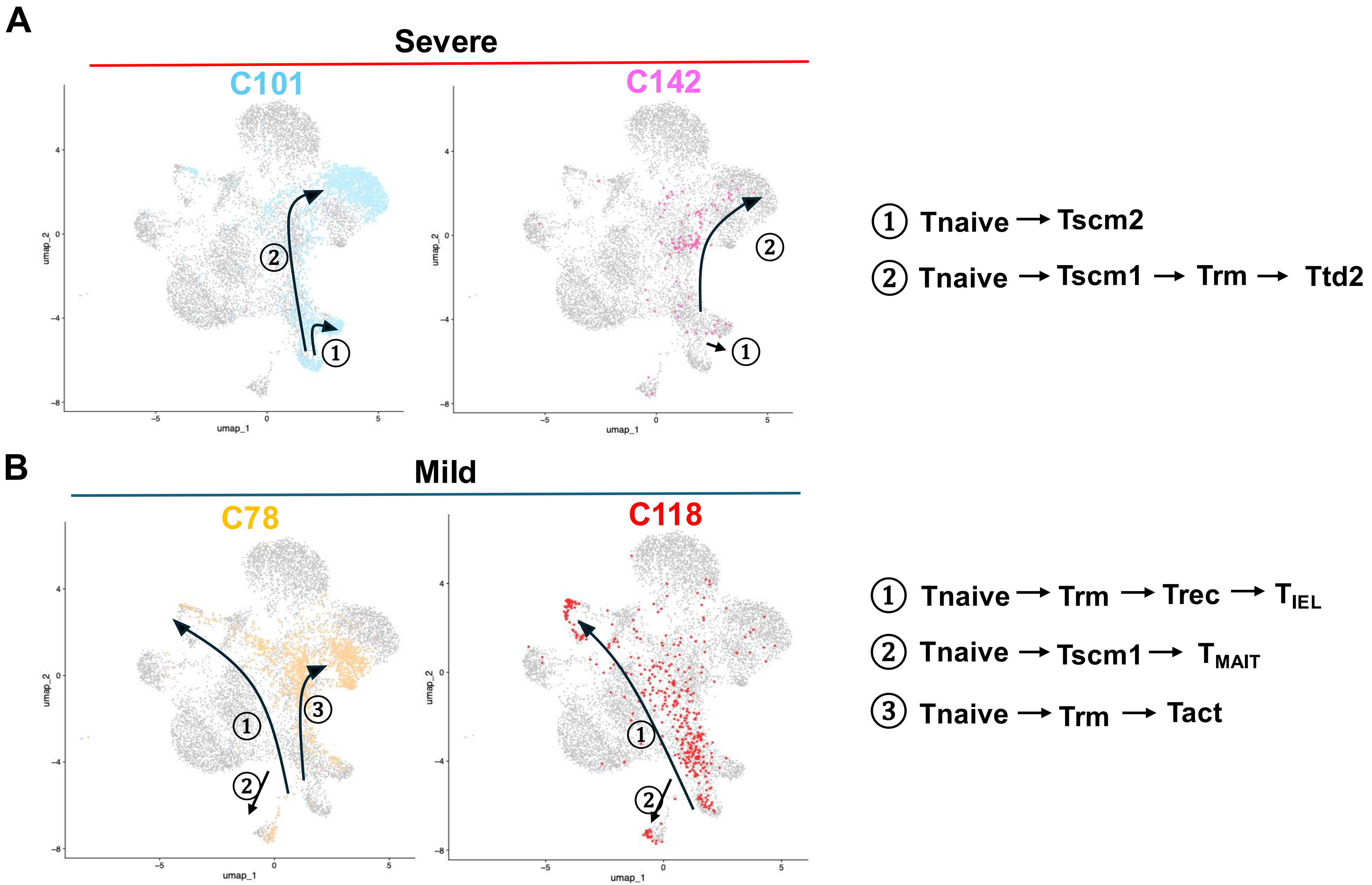

A
Severe
C101
C142
umap_1
umap_2
① Tnaive → Tscm2
② Tnaive → Tscm1 → Trm → Ttd2
B
Mild
C78
C118
umap_1
umap_2
① Tnaive → Trm → Trec → TIEL
② Tnaive → Tscm1 → TMAIT
③ Tnaive → Trm → Tact

**Figure S6. Possible differentiation pathways of antigen-experienced HLA-A2 restricted CD8$^+$ T cells.** (A) Inferred differentiation patterns (① and ②) of antigen-experienced HLA-A2 restricted CD8$^+$ T cells in severe COVID-19 patients (C101 and C142). (B) Inferred differentiation patterns (①, ②, and ③) of antigen-experienced HLA-A2 restricted CD8$^+$ T cells in mild COVID-19 patients (C78 and C118).

**Table S1** Identification of 42 HLA-A2$^{+}$ COVID-19 patients out of 77 patients by genotyping or phenotyping

| Patient ID | Typing Method | HLA-A*02 |
|---|---|---|
| C0002 | FACS | Positive |
| C0006 | FACS | |
| C0008 | FACS | |
| C0009 | DNA | |
| C0011 | DNA | |
| C0017 | DNA | |
| C0018 | DNA | |
| C0022 | DNA | |
| C0028 | DNA | |
| C0029 | DNA | |
| C0030 | DNA | |
| C0032 | DNA | |
| C0034 | FACS | |
| C0035 | FACS | |
| C0039 | DNA | |
| C0041 | DNA | |
| C0042 | DNA | |
| C0043 | DNA | |
| C0044 | FACS | |
| C0046 | FACS | |
| C0047 | DNA | |

| Patient ID | Typing Method | HLA-A*02 |
|---|---|---|
| C0051 | DNA | Positive |
| C0052 | DNA | |
| C0054 | DNA | |
| C0056 | FACS | |
| C0069 | FACS | |
| C0083 | DNA | |
| C0084 | FACS | |
| C0086 | FACS | |
| C0088 | DNA | |
| C0094 | DNA | |
| C0098 | DNA | |
| C0101 | DNA | |
| C0109 | FACS | |
| C0112 | FACS | |
| C0115 | FACS | |
| C0118 | DNA | |
| C0126 | FACS | |
| C0129 | DNA | |
| C0138 | DNA | |
| C0141 | DNA | |
| C0142 | DNA | |

**Table S2 Grading of the severity of COVID-19 patients at CUIMC**

1. I have not sought medical care
2. I went to an outpatient doctor but was not tested
3. I went to an outpatient doctor and was tested
4. I went to the Emergency Room and was not tested
5. I went to the Emergency room and was tested
6. I was admitted to the hospital but did not require extra oxygen
7. I was admitted to the hospital and did require extra oxygen
8. I was admitted to the hospital and went to the ICU
9. I was admitted to the hospital and required intubation (a tube to help you breathe)
10. I was admitted to the hospital and required ECMO (a heart and lung machine)

If a participant chooses between 1 and 5, they were noted as Mild.
If a participant chooses between 5 and 7, they were noted as Moderate
If a participant chooses between 8 and 10, they were noted as Severe

**Table S3**

### Predicted HLA-A2-binding peptides from WT SARS-CoV-2 AA sequence

| Name of the protein | ORF | No. | AA sequence | AA position | Ref_Accession_number | A2-binding Score (Percentile Rank) | Hydrophobicity |
|---|---|---|---|---|---|---|---|
| ExoN | ORF1ab polyprotein | 1 | YVNKHAFHT | 416-424 | YP_009724389.1 | 7.5 | 33.33% (Basic) |
| RdR Poly | ORF1ab polyprotein | 2 | VLTLDNQDL | 204-212 | YP_009724389.1 | 9.6 | 44.44% (Acid) |
| Helicase | ORF1ab polyprotein | 3 | AQLPAPRTL | 403-411 | YP_009724389.1 | 8.4 | 66.67% (Basic) |
| ExoN | ORF1ab polyprotein | 4 | IVCRFDTRV | 397-405 | YP_009724389.1 | 8.7 | 44.44% (Acid/Basic) |
| Helicase | ORF1ab polyprotein | 5 | FLCCKCCYD | 24-32 | YP_009724389.1 | 6.8 | 22.22% (Acid/Basic) |
| RdR Poly | ORF1ab polyprotein | 6 | FAQDGNAAI | 442-450 | YP_009724389.1 | 4.5 | 55.56% (Acid) |
| RdR Poly | ORF1ab polyprotein | 7 | TQMNLKYAI | 540-548 | YP_009724389.1 | 4 | 44.44% (Basic) |
| RdR Poly | ORF1ab polyprotein | 8 | HLMGWDYPK | 613-621 | YP_009724389.1 | 4 | 44.44% (Acid/Basic) |
| Helicase | ORF1ab polyprotein | 9 | YTACSHAAV | 306-314 | YP_009724389.1 | 2.6 | 44.44% (Basic) |
| ExoN | ORF1ab polyprotein | 10 | LQLGFSTGV | 107-115 | YP_009724389.1 | 1.3 | 44.44% (Neut) |
| Nsp7 | ORF1ab polyprotein | 11 | VLLSVLQQL | 12-20 | YP_009724389.1 | 1.3 | 66.67% (Neut) |
| RdR Poly | ORF1ab polyprotein | 12 | RLANECAQV | 654-662 | YP_009724389.1 | 1.1 | 44.44% (Acid/Basic) |
| RdR Poly | ORF1ab polyprotein | 13 | SLAIDAYPL | 861-869 | YP_009724389.1 | 0.8 | 66.67% (Acid) |
| 3CL Proteinase | ORF1ab polyprotein | 14 | FLNRFTTTL | 219-227 | YP_009724389.1 | 0.6 | 44.44% (Basic) |
| 3CL Proteinase | ORF1ab polyprotein | 15 | FLNGSCGSV | 140-148 | YP_009724389.1 | 0.6 | 33.33% (Neu) |
| PLpro | ORF1ab polyprotein | 16 | FLGRYMSAL | 77-85 | YP_009724389.1 | 0.6 | 55.56% (Basic) |
| SUD-M | ORF1ab polyprotein | 17 | ILGTVSWNL | 0-8 | YP_009724389.1 | 1.2 | 55.56% (Neu) |
| RdRp | ORF1ab polyprotein | 18 | TMADLVYAL | 123-131 | YP_009724389.1 | 0.3 | 66.67% (Acid) |
| RdRp | ORF1ab polyprotein | 19 | LMIERFVSL | 854-862 | YP_009724389.1 | 0.5 | 66.67% (Acid/Basic) |
| RdRp | ORF1ab polyprotein | 20 | ILHCANFNV | 307-315 | YP_009724389.1 | 0.6 | 55.56% (Basic) |
| RdRp | ORF1ab polyprotein | 21 | YTMADLVYA | 122-130 | YP_009724389.1 | 0.7 | 55.56% (Acid) |
| RdRp | ORF1ab polyprotein | 22 | RLSFKELLV | 365-373 | YP_009724389.1 | 1.9 | 55.56% (Acid/Basic) |
| S protein | Spike glycoprotein | 23 | VLNDILSRL | 976-984 | QHR63260.2 | 1.1 | 55.56% (Acid/Basic) |
| S protein | Spike glycoprotein | 24 | VVFLHVTYV | 1060-1068 | QHR63260.2 | 1.2 | 66.67% (Basic) |
| N protein | Nucleocapsid protein | 25 | LLLDRLNQL | 222-230 | WBP61198.1 | 0.8 | 55.56% (Acid/Basic) |
| Helicase | ORF1ab polyprotein | 26 | KLSYGIATV | 146-154 | YP_009724389.1 | 0.4 | 44.44 (Basic) |
| Helicase | ORF1ab polyprotein | 27 | KLNVGDYFV | 218-226 | YP_009724389.1 | 0.5 | 44.44 (Acid/Basic) |
| Helicase | ORF1ab polyprotein | 28 | KLQFTSLEI | 584-592 | YP_009724389.1 | 2.3 | 44.44 (Acid/Basic) |
| Helicase | ORF1ab polyprotein | 29 | IVDTVSALV | 448-456 | YP_009724389.1 | 2.4 | 66.67 (Acid) |
| S protein | Spike glycoprotein | 30 | NLNESLIDL | 1192-1200 | QHR63260.2 | 2.8 | 44.44% (Acid) |
| S protein | Spike glycoprotein | 31 | RLNEVAKNL | 1185-1193 | QHR63260.2 | 5.1 | 44.44% (Acid/Basic) |

**Table S4 HLA-A2-restricted CD8$^{+}$ T-cell responses of 24 COVID patients to specific epitopes of SARS-CoV-2, as determined by ELISpot assay**

| Name of the protein | No. | AA sequence | prevalence score (# of patient responded) | C0002 (15d) | C0006 (22d) | C0006 (62d) | C0008 (91d) | C0009 (119d) | C0017 (39d) | C0017 (121d) | C0018 (38d) | C0018 (113d) | C0028 (130d) | C0030 (90d) | C0032 (35d) | C0032 (116d) | C0034 (26d) | C0035 (39d) | C0042 (43d) | C0044 (59d) | C0046 (50d) | C0051 (38d) | C0051 (104d) | C0056 (47d) | C0069 (61d) | C0078 (32d) | C0086 (25d) | C0086 (31d) | C0109 (4d) | C0109 (18d) | C0115 (38d) | C0118 (66d) | C0126 (9d) | C0141 (98d) |
|---|---|---|---|---|---|---|---|---|---|---|---|---|---|---|---|---|---|---|---|---|---|---|---|---|---|---|---|---|---|---|---|---|---|---|
| ExoN | 1 | YVNKHAFHT | 1 (2) | | | | | | | | | | | | | | | +/- | | | | | | | +/- | | | | | | | | | |
| RdR Poly | 2 | VLTLDNQDL | 1.5 (2) | | | | | | | | | | | | | | | +/- | | | + | | | | | | | | | | | | | |
| Helicase | 3 | AQLPAPRTL | 1.5 (2) | + | | | | | | | | | | | | | | +/- | | | | | | | | | | | | | | | | |
| ExoN | 4 | IVCRFDTRV | 2.5 (3) | | | | +/- | | | | | | | | | | | | | | | | | | + | | | | | | + | | | |
| Helicase | 5 | FLCCKCCYD | 0.5 (1) | | | | | | | | | | | | | | | +/- | | | | | | | | | | | | | | | | |
| RdR Poly | 6 | FAQDGNAAI | 6 (6) | | | | +/- | | | | | | + | | | | | +/- | | | | | + | | | | ++ | + | | | | | | |
| RdR Poly | 7 | TQMNLKYAI | 2 (4) | +/- | | | | | | | | | | | | | +/- | | | | | | | | + | | | | | | +/- | | | |
| RdR Poly | 8 | HLMGWDYPK | 2 (1) | | | | ++ | | | | | | | | | | | | | | | | | | | | | | | | | | | |
| Helicase | 9 | YTACSHAAV | 0 | | | | | | | | | | | | | | | | | | | | | | | | | | | | | | | |
| ExoN | 10 | LQLGFSTGV | 1 (1) | | | | | | | | | | | | | | | + | | | | | | | | | | | | | | | | |
| Nsp7 | 11 | VLLSVLQQL | 1 (2) | | | | | | | | | | | | | | | | | | +/- | | | | | | +/- | | | | | | | |
| RdR Poly | 12 | RLANECAQV | 1.5 (2) | | | | | | | | | | | | | | | | | + | | | | | | | | | | | | | +/- | |
| RdR Poly | 13 | SLAIDAYPL | 2 (2) | | | | | | | | | | | | | | | | | | + | | | | | | | | | | + | | | |
| 3CL Proteinase | 14 | FLNRFTTTL | 0.5 (1) | | | | | | | | | | | | | | | | | | | | | +/- | | | | | | | | | | |
| 3CL Proteinase | 15 | FLNGSCGSV | 2 (1) | | | | | | | | | | | | | | | | | | | | | | | | ++ | | | | | | | |
| PLpro | 16 | FLGRYMSAL | 11 (11) | ++ | | | | | | ++ | + | + | | | | | | +/- | | | | + | + | +/- | +/- | | | | +/- | | | + | | |
| SUD-M | 17 | ILGTVSWNL | 11.5 (10) | | | | +/- | | | + | ++ | | | | | | | +/- | | + | | | ++ | | | ++ | +/- | | + | | | + | | |
| RdRp | 18 | TMADLVYAL | 1 (2) | | | | +/- | | | | | | | | | | | +/- | | | | | | | | | | | | | | | | |
| RdRp | 19 | LMIERFVSL | 23 (15) | +++ | ++ | +++ | | + | + | | | | | + | | | +/- | +/- | ++ | | | | ++ | | + | ++ | | | | | | ++ | + | + |
| RdRp | 20 | ILHCANFNV | 9 (7) | | | | ++ | | | | + | | | | | | ++ | +/- | | | | + | ++ | | +/- | | | | | | | | | |
| RdRp | 21 | YTMADLVYA | 5.5 (5) | | | | | | | | + | | | | | + | | +/- | | | | | | | | | ++ | | | | + | | | |
| RdRp | 22 | RLSFKELLV | 1.5 (2) | | | | +/- | | | | | | | | | | | | | | | | | | | | + | | | | | | | |
| S protein | 23 | VLNDILSRL | 6 (5) | | | | | | | | + | | | | | | ++ | +/- | | | | | | | | | ++ | | | | | | +/- | |
| S protein | 24 | VVFLHVTYV | 22.5 (18) | | | | | | | | | ++ | + | ++ | + | | | ++ | +++ | + | +/- | | + | + | + | | + | +/- | + | + | +/- | | ++ | + |
| N protein | 25 | LLLDRLNQL | 2 (3) | | | | + | | | | | | | | | | | | | | | | | +/- | | | | | | | | | +/- | |
| Helicase | 26 | KLSYGIATV | 6.5 (6) | +/- | | | | | | | + | | + | | | | + | + | | | | | | | ++ | | | | | | | | | |
| Helicase | 27 | KLNVGDYFV | 1.5 (2) | | | | | | | | | | | | | | | | | | +/- | | | | | | | | | | + | | | |
| Helicase | 28 | KLQFTSLEI | 1.5 (2) | | | | | | | | | | | | | | | + | | | +/- | | | | | | | | | | | | | |
| Helicase | 29 | IVDTVSALV | 1 (1) | | | | | | | | | | | | | | | | | + | | | | | | | | | | | | | | |
| S protein | 30 | NLNESLIDL | 4 (4) | | | | + | | | | | | | | | | | | | | +/- | | | | | | ++ | | | | +/- | | | |
| S protein | 31 | RLNEVAKNL | 2 (2) | | | | + | | | | | | | | | | | | | | | | | | | | | | | | + | | | |

++; stimulation index= >2, +; SI= 2>1.5, +/-; SI= 1.5>1.2

### Table S5 Sequence homology of SARS-CoV-2 epitopes among SARS-CoV-2 variants and other strains

| | AA Sequence | SARS1 | Alpha | Beta | Gamma | Delta | BA.1 | BA.5 | XBB.1.5 | EG.5 | JN.1 | KP.2 | KP3.1.1 | KP.3 (LB.1 and KP2.3) | XEC | MERS | NL6 | 229E | HKU |
|---|---|---|---|---|---|---|---|---|---|---|---|---|---|---|---|---|---|---|---|
| ExoN | IVCRFDTRV | Yes (100%); No (0%); Total: 133 | Yes (97%); No (3%); Total: 200 | Yes (97.5%); No (2.5%); Total: 200 | Yes (97.5%); No (2.5%); Total: 200 | Yes (94.3%); No (5.7%); Total: 193 | Yes (97.5%); No (2.5%); Total: 200 | Yes (98%); No (2%); Total: 200 | Yes (98%); No (2%); Total: 200 | Yes (97%); No (3%); Total: 200 | Yes (100%); No (0%); Total: 200 | Yes (98.5%); No (1.5%); Total: 200 | Yes (99.5%); No (0.5%); Total: 200 | Yes (98.5%); No (1.5%); Total: 200 | Yes (100%); No (0%); Total: 200 | Yes | No | No | Yes |
| Helicase | KLSYGIATV | Yes (100%); No (0%); Total: 133 | Yes (95.5%); No (4.5%); Total: 200 | Yes (95.5%); No (4.5%); Total: 200 | Yes (97.5%); No (2.5%); Total: 200 | Yes (86%); No (14%); Total: 193 | Yes (99.5%); No (0.5%); Total: 200 | Yes (99.5%); No (0.5%); Total: 200 | Yes (98.5%); No (1.5%); Total: 200 | Yes (98.5%); No (1.5%); Total: 200 | Yes (99%); No (1%); Total: 200 | Yes (98.5%); No (1.5%); Total: 200 | Yes (100%); No (0%); Total: 200 | Yes (100%); No (0%); Total: 200 | Yes (99.5%); No (0.5%); Total: 200 | No | No | No | No |
| PLP | FLGRYMSAL | Yes (100%); No (0%); Total: 133 | Yes (93.5%); No (6.5%); Total: 200 | Yes (93%); No (7%); Total: 200 | Yes (96%); No (4%); Total: 200 | Yes (77.2%); No (22.8%); Total: 193 | Yes (96%); No (4%); Total: 200 | Yes (94.5%); No (5.5%); Total: 200 | Yes (93%); No (7%); Total: 200 | Yes (97.5%); No (2.5%); Total: 200 | Yes (98.5%); No (1.5%); Total: 200 | Yes (95.5%); No (4.5%); Total: 200 | Yes (98.5%); No (1.5%); Total: 200 | Yes (97.5%); No (2.5%); Total: 200 | Yes (97.5%); No (2.5%); Total: 200 | No | No | No | No |
| RdR Poly | YTMADLVYA | Yes (98.5%); No (1.5%); Total: 133 | Yes (99%); No (1%); Total: 200 | Yes (97%); No (3%); Total: 200 | Yes (99.5%); No (0.5%); Total: 200 | Yes (99%); No (1%); Total: 193 | Yes (99.5%); No (0.5%); Total: 200 | Yes (97%); No (3%); Total: 200 | Yes (98.5%); No (1.5%); Total: 200 | Yes (98.5%); No (1.5%); Total: 200 | Yes (99.5%); No (0.5%); Total: 200 | Yes (99.5%); No (0.5%); Total: 200 | Yes (100%); No (0%); Total: 200 | Yes (100%); No (0%); Total: 200 | Yes (99.5%); No (0.5%); Total: 200 | No | No | No | No |
| RdRp1 | ILHCANFNV | Yes (97.7%); No (2.3%); Total: 133 | Yes (96.5%); No (3.5%); Total: 200 | Yes (98%); No (2%); Total: 200 | Yes (99%); No (1%); Total: 200 | Yes (92.2%); No (7.8%); Total: 193 | Yes (98.5%); No (1.5%); Total: 200 | Yes (97%); No (3%); Total: 200 | Yes (96.5%); No (3.5%); Total: 200 | Yes (95.5%); No (4.5%); Total: 200 | Yes (99.5%); No (0.5%); Total: 200 | Yes (98%); No (2%); Total: 200 | Yes (100%); No (0%); Total: 200 | Yes (100%); No (0%); Total: 200 | Yes (100%); No (0%); Total: 200 | No | No | No | No |
| RdRp2 | FAQDGNAAI | Yes (100%); No (0%); Total: 133 | Yes (98.5%); No (1.5%); Total: 200 | Yes (97.5%); No (2.5%); Total: 200 | Yes (99.5%); No (0.5%); Total: 200 | Yes (93.8%); No (6.2%); Total: 193 | Yes (97%); No (3%); Total: 200 | Yes (92%); No (8%); Total: 200 | Yes (95%); No (5%); Total: 200 | Yes (96.5%); No (3.5%); Total: 200 | Yes (99%); No (1%); Total: 200 | Yes (98%); No (2%); Total: 200 | Yes (98%); No (2%); Total: 200 | Yes (97.5%); No (2.5%); Total: 200 | Yes (1000%); No (0%); Total: 200 | Yes | No | No | No |
| RdRp3 | LMIERFVSL | Yes (100%); No (0%); Total: 133 | Yes (98.5%); No (1.5%); Total: 200 | Yes (98.5%); No (1.5%); Total: 200 | Yes (99.5%); No (0.5%); Total: 200 | Yes (89.6%); No (10.4%); Total: 193 | Yes (99%); No (1%); Total: 200 | Yes (99%); No (1%); Total: 200 | Yes (98%); No (2%); Total: 200 | Yes (94%); No (6%); Total: 200 | Yes (98.5%); No (1.5%); Total: 200 | Yes (98%); No (2%); Total: 200 | Yes (99%); No (1%); Total: 200 | Yes (98.5%); No (1.5%); Total: 200 | Yes (98.5%); No (1.5%); Total: 200 | No | No | No | No |
| S protein1 | VLNDILSRL | Yes (94%); No (6%); Total: 133 | Yes (2%); No (98%); Total: 200 | Yes (96%); No (4%); Total: 200 | Yes (98%); No (2%); Total: 200 | Yes (88%); No (12%); Total: 200 | Yes (1%); No (99%); Total: 200 | Yes (96%); No (4%); Total: 200 | Yes (93%); No (7%); Total: 200 | Yes (94.5%); No (5.5%); Total: 200 | Yes (77%); No (23%); Total: 200 | Yes (94%); No (6%); Total: 200 | Yes (96.5%); No (3.5%); Total: 200 | Yes (95.5%); No (4.5%); Total: 200 | Yes (98%); No (2%); Total: 200 | No | No | No | No |
| S protein2 | VVFLHVTYV | Yes (97.7%); No (2.3%); Total: 133 | Yes (97%); No (3%); Total: 200 | Yes (96%); No (4%); Total: 200 | Yes (99%); No (1%); Total: 200 | Yes (94%); No (6%); Total: 200 | Yes (92%); No (8%); Total: 200 | Yes (98.5%); No (1.5%); Total: 200 | Yes (93%); No (7%); Total: 200 | Yes (96%); No (4%); Total: 200 | Yes (77.5%); No (22.5%); Total: 200 | Yes (93%); No (7%); Total: 200 | Yes (94.5%); No (5.5%); Total: 200 | Yes (95.5%); No (4.5%); Total: 200 | Yes (97%); No (3%); Total: 200 | No | No | No | No |
| S protein3 | NLNESLIDL | Yes (98.5%); No (1.5%); Total: 133 | Yes (96%); No (4%); Total: 200 | Yes (93.5%); No (6.5%); Total: 200 | Yes (98.5%); No (1.5%); Total: 200 | Yes (92.5%); No (7.5%); Total: 200 | Yes (87.5%); No (12.5%); Total: 200 | Yes (97%); No (3%); Total: 200 | Yes (91%); No (9%); Total: 200 | Yes (96%); No (4%); Total: 200 | Yes (74.5%); No (25.5%); Total: 200 | Yes (90.5%); No (9.5%); Total: 200 | Yes (96%); No (4%); Total: 200 | Yes (91.5%); No (8.5%); Total: 200 | Yes (95%); No (5%); Total: 200 | No | No | No | No |
| SUD-M | ILGTVSWNL | Yes (100%); No (0%); Total: 133 | Yes (98.5%); No (1.5%); Total: 200 | Yes (98%); No (2%); Total: 200 | Yes (97.50%); No (2.5%); Total: 200 | Yes (95.9%); No (4.1%); Total: 193 | Yes (97%); No (3%); Total: 200 | Yes (89.5%); No (10.5%); Total: 200 | Yes (87.5%); No (12.5%); Total: 200 | Yes (87%); No (13%); Total: 200 | Yes (93.5%); No (6.5%); Total: 200 | Yes (96.5%); No (3.5%); Total: 200 | Yes (100%); No (0%); Total: 200 | Yes (99.5%); No (0.5%); Total: 200 | Yes (91%); No (9%); Total: 200 | No | No | No | No |

### Table S6 Cell hashtag list

| Patient # | Hash # | Product | Catalog # | Barcode seq | Single cell data |
|---|---|---|---|---|---|
| C29 (Mild COVID-19 patient) | Hash3 | TotalSeq™-B0253 anti-human Hashtag 3 Antibody | 394635 | TTCCGCCTCTCTTTG | MKH |
| C30 (Mild COVID-19 patient) | Hash4 | TotalSeq™-B0254 anti-human Hashtag 4 Antibody | 394637 | AGTAAGTTCAGCGTA | |
| C32 (Mild COVID-19 patient) | Hash5 | TotalSeq™-B0255 anti-human Hashtag 5 Antibody | 394639 | AAGTATCGTTTCGCA | |
| C44 (Severe COVID-19 patient) | Hash6 | TotalSeq™-B0256 anti-human Hashtag 6 Antibody | 394641 | GGTTGCCAGATGTCA | |
| C52 (Severe COVID-19 patient) | Hash7 | TotalSeq™-B0257 anti-human Hashtag 7 Antibody | 394643 | TGTCTTTCCTGCCAG | |
| C78 (Mild COVID-19 patient) | Hash2 | TotalSeq™-B0252 anti-human Hashtag 2 Antibody | 394633 | TGATGGCCTATTGGG | MKK |
| C86 (Mild COVID-19 patient) | Hash3 | TotalSeq™-B0253 anti-human Hashtag 3 Antibody | 394635 | TTCCGCCTCTCTTTG | |
| C101 (Severe COVID-19 patient) | Hash4 | TotalSeq™-B0254 anti-human Hashtag 4 Antibody | 394637 | AGTAAGTTCAGCGTA | |
| C118 (Mild COVID-19 patient) | Hash5 | TotalSeq™-B0255 anti-human Hashtag 5 Antibody | 394639 | AAGTATCGTTTCGCA | |
| C138 (Severe COVID-19 patient) | Hash6 | TotalSeq™-B0256 anti-human Hashtag 6 Antibody | 394641 | GGTTGCCAGATGTCA | |
| C142 (Severe COVID-19 patient) | Hash7 | TotalSeq™-B0257 anti-human Hashtag 7 Antibody | 394643 | TGTCTTTCCTGCCAG | |

#### Table S7 CITE-seq antibody list

| Cat# | Product name | Marker | Antibody Barcode |
|---|---|---|---|
| 353249 | TotalSeq™-B0148 anti-human CD197 (CCR7) Antibody | CCR7 | AGTTCAGTCAACCGA |
| 304161 | TotalSeq™-B0063 anti-human CD45RA Antibody | CD45RA | TCAATCCTTCCGCTT |
| 350235 | TotalSeq™-B0145 anti-human CD103 (Integrin αE) Antibody | CD103 | GACCTCATTGTGAAT |
| 103071 | TotalSeq™-B0125 anti-human CD44 Antibody | CD44 | AATCCTTCCGAATGT |
| 367727 | TotalSeq™-B0153 anti-human KLRG1 (MAFA) Antibody | KLRG1 | CTTATTTCCTGCCCT |
| 329961 | TotalSeq™-B0088 anti-human CD279 (PD-1) Antibody | PD-1 | ACAGCGCCGTATTTA |
| 369629 | TotalSeq™-B0151 anti-human CD152 (CTLA-4) Antibody | CTLA-4 | ATGGTTCACGTAATC |
| 345053 | TotalSeq™-B0169 anti-human CD366 (Tim-3) Antibody | TIM-3 | TGTCCTACCCAACTT |
| 356639 | TotalSeq™-B0410 anti-human CD38 Antibody | CD38 | CCTATTCCGATTCCG |
| 307661 | TotalSeq™-B0159 anti-human HLA-DR Antibody | HLA-DR | AATAGCGAGCAAGTA |
| 369337 | TotalSeq™-B0152 anti-human CD223 (LAG-3) Antibody | LAG3 | CATTTGTCTGCCGGT |
| 309837 | TotalSeq™-B0355 anti-human CD137 (4-1BB) Antibody | CD137 | CAGTAAGTTCGGGAC |
| 310949 | TotalSeq™-B0146 anti-human CD69 Antibody | CD69 | GTCTCTTGGCTTAAA |
| 304347 | TotalSeq™-B0576 anti-human CD49d Antibody | CD49d | CCATTCAACTTCCGG |
| 302647 | TotalSeq™-B0085 anti-human CD25 Antibody | CD25 | TTTGTCCTGTACGCC |
| 320839 | TotalSeq™-B0165 anti-human CD314 (NKG2D) Antibody | NKG2D | CGTGTTTGTTCCTCA |

**Table S8** **List of top 30 genes in $T_{IEL\text{-}like}$ and C1 cluster by differentially expressed gene (DEG) analysis, respectively**

| Tiel-like cluster (top 30 genes) | C1 cluster (top 30 genes) |
|---|---|
| TYROBP | CHEK1 |
| TRDC | TYMS |
| KLRF1 | CDT1 |
| CEBPD | ID3 |
| FCGR3A | RBBP8 |
| XAF1 | CLSPN |
| CD300A | CTNNAL1 |
| SH2D1B | TNFRSF9 |
| EPSTI1 | HLA-DRB5 |
| IFI44L | LTB |
| PRF1 | COPRS |
| FCER1G | RRM2 |
| CLIC3 | HIST1H2AL |
| IFI6 | FAM111B |
| MX1 | TOP2A |
| GNLY | BIRC5 |
| KLRC3 | UBE2T |
| IL2RB | MTHFD2 |
| SAMD9L | CENPW |
| OAS1 | FABP5 |
| CD300A | NUSAP1 |
| GNPTAB | NME1 |
| MX2 | ASPM |
| EIF2AK2 | SPC25 |
| IFIT1 | HMMR |
| HIPK2 | KNL1 |
| HERC5 | TPX2 |
| IFITM1 | UBE2C |
| KLRD1 | GTSE1 |
| CTSD | CENPU |
